# Decoration of the enterococcal polysaccharide antigen EPA is essential for virulence, cell surface charge and resistance to innate immunity

**DOI:** 10.1101/479022

**Authors:** Robert E. Smith, Bartłomiej Salamaga, Piotr Szkuta, Natalia Hajdamowicz, Tomasz K. Prajsnar, Gregory Bulmer, Thierry Fontaine, Justyna Kołodziejczyk, Jean-Marie Herry, Andrea Hounslow, Mike P. Williamson, Pascale Serror, Stéphane Mesnage

**Affiliations:** Krebs Institute, University of Sheffield, Firth Court, Western Bank, Sheffield S10 2TN, UK; Department of Molecular Biology and Biotechnology, University of Sheffield, Firth Court, Western Bank, Sheffield S10 2TN, UK; Unité des Aspergillus, Institut Pasteur, 75015 Paris, France; Micalis Institute, INRA, AgroParisTech, Université Paris-Saclay, 78350 Jouy en Josas, France

## Abstract

*Enterococcus faecalis* is an opportunistic pathogen with an intrinsically high resistance to lysozyme, a key effector of the innate immune system. This high level of resistance requires several genes (*oatA, pgdA, dltA* and *sigV*) acting synergistically to inhibit both the enzymatic and cationic antimicrobial peptide activities of lysozyme. We sought to identify novel genes modulating *E. faecalis* resistance to lysozyme. Random transposon mutagenesis carried out in the quadruple *oatA*/*pgdA*/*dltA*/*sigV* mutant led to the identification of several independent insertions clustered on the chromosome. These mutations were located in a locus referred to as the enterococcal polysaccharide antigen (EPA) variable region located downstream of the highly conserved *epaA-epaR* genes proposed to encode a core synthetic machinery. The *epa* variable region was previously proposed to be responsible for EPA decorations, but the role of this locus remains largely unknown. Here, we show that EPA decoration contributes to resistance towards charged antimicrobials and underpins virulence in the zebrafish model of infection by conferring resistance to phagocytosis. Collectively, our results indicate that the production of the EPA rhamnopolysaccharide backbone is not sufficient to promote *E. faecalis* infections and reveal an essential role of the modification of this surface polymer for enterococcal pathogenesis.

## Introduction

*Enterococcus faecalis* is a commensal bacterium found in the gastro-intestinal tract of humans and frequently isolated from the environment as a result of faecal contamination [1, 2]. Although this organism is considered as harmless in healthy carriers, *E. faecalis* has been proposed to contribute to the pathogenesis of inflammatory bowel disease and colorectal cancer [3, 4]. *E. faecalis* can also cause a wide range of hospital-acquired opportunistic infections that can be life-threatening [1]. *E. faecalis* infections can be difficult to treat due to the resistance of this organism to antibiotics such as cephalosporins and glycopeptides (Vancomycin Resistant Enterococci, VRE) and its capacity to form biofilms [5]. Interestingly, *E. faecalis* strains responsible for hospital-acquired infections are also found in healthy individuals and genes associated with virulence are not only present in clinical isolates [6]. How this organism can cause infections is therefore not entirely understood. One property of *E. faecalis* that contributes to the onset of infections is its resistance to the host innate immune system. Cell surface polymers including teichoic acids (TAs), a capsule and the enterococcal polysaccharide antigen (EPA) confer phagocytosis evasion and resistance to complement activation [7-9]. *E. faecalis* also displays an intrinsically high resistance to lysozyme, a key component of the innate immune system representing a first line of defence against pathogens. Lysozyme is found in virtually all human biological fluids including saliva, milk, serum and tears where it is found at concentrations between 1-2 mg ml^−1^ [10, 11]. Lysozyme has two distinct antimicrobial activities. Firstly, it hydrolyses the glycan chains of peptidoglycan, the major component of the bacterial cell wall, causing cell lysis [12]. Secondly, lysozyme displays cationic antimicrobial peptide (CAMP) activity. Lysozyme contains highly charged C-terminal sequences (RAWVAWRNR in human lysozyme) sufficient to inhibit bacterial growth [13] by causing membrane permeabilization [14].

Four genes (*oatA, pgdA, dltA* and *sigV*) contribute synergistically to lysozyme resistance in *E. faecalis*. Both OatA and PgdA modify peptidoglycan glycan strands, thereby inhibiting lysozyme catalytic activity. OatA is an *O*-acetyl transferase that transfers an acetyl group onto the C6-OH group of *N*-acetylmuramic acid residues [15]. PgdA is produced in response to lysozyme and is an esterase that removes the acetyl group in position 2 of *N*-acetylglucosamine residues [16]. DltA is a D-alanine-D-alanyl carrier ligase essential for the alanylation of TAs. It has been proposed that this modification reduces the net negative charge of TAs and inhibits the CAMP activity of lysozyme [17]. SigV is an extracytoplasmic sigma factor that controls the expression level of *pgdA* in response to lysozyme [18, 19]. *oatA, pgdA, dltA* and *sigV* act in concert to confer high-level resistance to lysozyme in *E. faecalis.* Deletions in these genes (alone or in combination) have been associated with a reduction in virulence in mice or *Galleria mellonella* [18, 19] and a decrease in survival within murine peritoneal macrophages [15].

In this study, we show that the quadruple mutant (*oatA, pgdA, dltA* and *sigV*; *OPDV* strain) still displays a relatively high resistance to lysozyme in comparison to other Firmicutes. Using transposon mutagenesis, we used this quadruple mutant to carry out transposon mutagenesis and to identify additional genes involved in lysozyme resistance. We show that several genes contributing to the decoration of the enterococcal polysaccharide antigen play an essential role in the resistance to effectors of the innate immune system and in virulence.

## Results

### The *E. faecalis* quadruple mutant harboring deletions in *oatA, pgdA, dltA* and *sigV* displays a relatively high residual resistance to lysozyme

We determined the minimal inhibitory concentrations (MIC) of lysozyme for several Gram-positive bacteria (Table 1; see S1 Fig for a representative set of MIC assays). *Micrococcus luteus*, used as a reference substrate to define lysozyme activity [20] had an expected very low MIC of 5 × 10^−4^ mg ml^−1^. MIC values were higher for all Firmicutes tested. Growth of several species was inhibited by lysozyme concentrations of 0.0312 mg ml^−1^ (*Aerococcus viridans Bacillus subtilis, Bacillus megaterium* and *Lactobacillus cellobiosus*). *Lactococcus lactis* growth was inhibited by concentrations of 0.125 mg ml^−1^. The MIC of lysozyme for all pathogens tested was relatively high: 4 mg ml^−1^ for *Listeria monocytogenes* and >16 mg ml^−1^ for *Staphylococcus aureus,* and all enterococci and streptococci tested. In *L. monocytogenes*, lysozyme resistance was largely due to peptidoglycan de-*N*-acetylation. Deletion of the gene encoding the deacetylase PgdA led to a 32-fold decrease in resistance (MIC=0.125 mg ml^−1^). Interestingly, abolishing peptidoglycan *O*-acetylation or de-*N*-acetylation in *E. faecalis* only had a minor impact on lysozyme resistance [16, 19]. The combined deletions in the four genes contributing to *E. faecalis* lysozyme resistance (*oatA, pgdA, dltA* and *sigV; OPDV* strain) was required for 10-fold reduction in the MIC of this antimicrobial compound (0.5 mg ml^−1^). However, the lysozyme MIC for the *OPDV* mutant was still higher than that of non-pathogenic Gram-positive bacteria.

**Table 1.** MIC of lysozyme for Firmicutes.

| Strain | lysozyme MIC<br>(mg ml<sup>-1</sup>) |
| --- | --- |
| *Staphylococcus aureus* COL | >16 |
| *Streptococcus gallolyticus* UCN34 | >16 |
| *Streptococcus gordonii* DL-1 Challis | >16 |
| *Streptococcus mutans* UA159 | >16 |
| *Enterococcus faecium* DO | >16 |
| *Enterococcus hirae* ATCC9790 | >16 |
| *Enterococcus faecalis* OG1RF | >16 |
| *Enterococcus faecalis* O (ΔoatA) | >16 |
| *Enterococcus faecalis* OP (ΔoatA ΔpgdA) | 16 |
| *Enterococcus faecalis* OPD (ΔoatA ΔpgdA ΔdltA) | 8 |
| *Enterococcus faecalis* OPDV (ΔoatA ΔpgdA ΔdltA ΔsigV) | 0.5 |
| *Listeria monocytogenes* EGD | 4 |
| *Listeria monocytogenes* EGD ΔpgdA | 0.125 |
| *Lactococcus lactis* MG1363 | 0.125 |
| *Bacillus subtilis* 168 | 0.0312 |
| *Bacillus megaterium* KM | 0.0312 |
| *Lactobacillus cellobiosus* ATCC11739 | 0.0312 |
| *Aerococcus viridans* ATCC11563 | 0.0312 |
| *Micrococcus luteus* ATCC4698 | 0.0005 |

### Transposon mutagenesis of the *OPDV epa* variable region confers resistance to lysozyme

The relatively high lysozyme MIC of the quadruple mutant (*OPDV* strain) prompted us to further explore *E. faecalis* properties modulating lysozyme activity. We constructed a transposon mutant library in the *OPDV* background using the *Mariner*-based system previously described for *E. faecium* [21]. Transposon mutants were selected on agar plates containing lysozyme at a concentration of 2 mg ml^−1^, four times the MIC for the parental *OPDV* strain. Approximately 2 × 10^5^ mutants were plated and after 24 h incubation at 37°C, 16 mutants forming colonies at this concentration were isolated and further analysed. Mapping the transposon insertion sites revealed that 9 mutants had insertions downstream of the conserved *epaA-epaR* region encoding the core synthetic apparatus likely required to produce the enterococcal polysaccharide antigen EPA (Fig 1A) [22]. The region containing the insertions displays genetic variability between strains and was proposed to be responsible for the decoration of the EPA polysaccharide [22-24]. Mutations were clustered around three genes encoding putative glycosyltransferases and a homolog of *wcaG*, an epimerase/dehydratase (Fig 1B).

**Figure 1.**
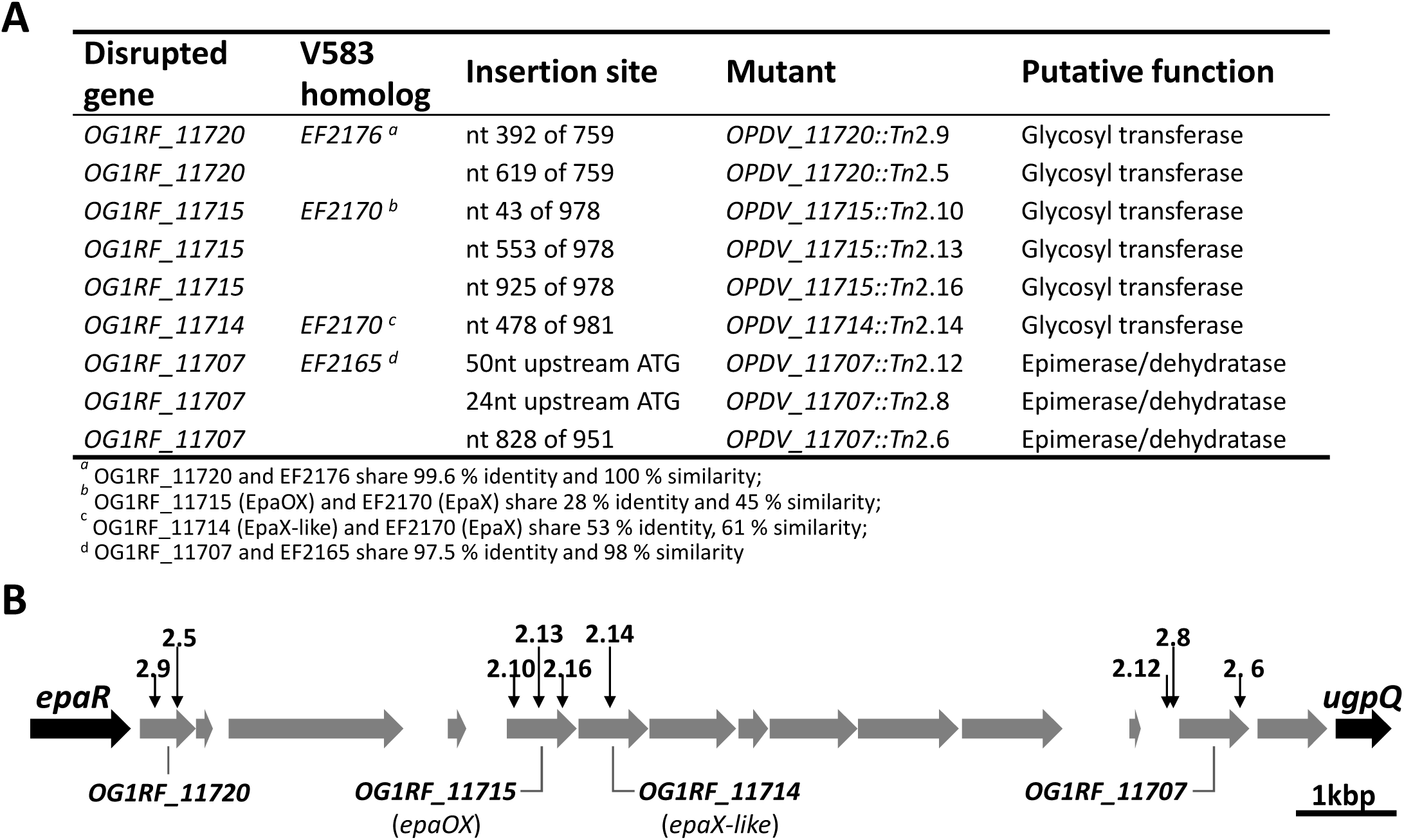
Identification of *epa* mutants resistant to lysozyme. **A.** Description of individual transposon insertions. **B.** Mapping of transposon insertions in the *epa* variable region. Insertion sites are indicated by vertical arrows. ORFs in the *epa* variable region are depicted in grey.

Transposon mutants were complemented to formally establish that the insertions in the *epa* variable region were responsible for lysozyme resistance. Four plasmids were constructed to express *OG1RF_11720, OG1RF_11715* (*epaOX*), *OG1RF_11714* (*epaX-like*) and *OG1RF_11707* under the control of the inducible *tet* promoter. Following transformation into *E. faecalis,* gene expression was induced in the presence of anhydrotetracycline (ATc) and the production of his-tagged proteins was checked by western blot (S2 Fig). Complementation was evaluated by measuring susceptibility to lysozyme (Fig 2). Lysozyme resistance associated with transposon insertions in genes *OG1RF_11720, OG1RF_11715, OG1RF_11714* and *OG1RF_11707* could be complemented when the disrupted gene was expressed in *trans*. By contrast, the parental susceptibility to lysozyme could not be restored when complementation experiments were carried out with plasmids encoding a gene distinct form the one disrupted (Table 2). Altogether, these experiments confirmed that the resistance phenotypes of the mutants analysed were due to the disruption of the genes indicated in Fig 1.

**Figure 2.**
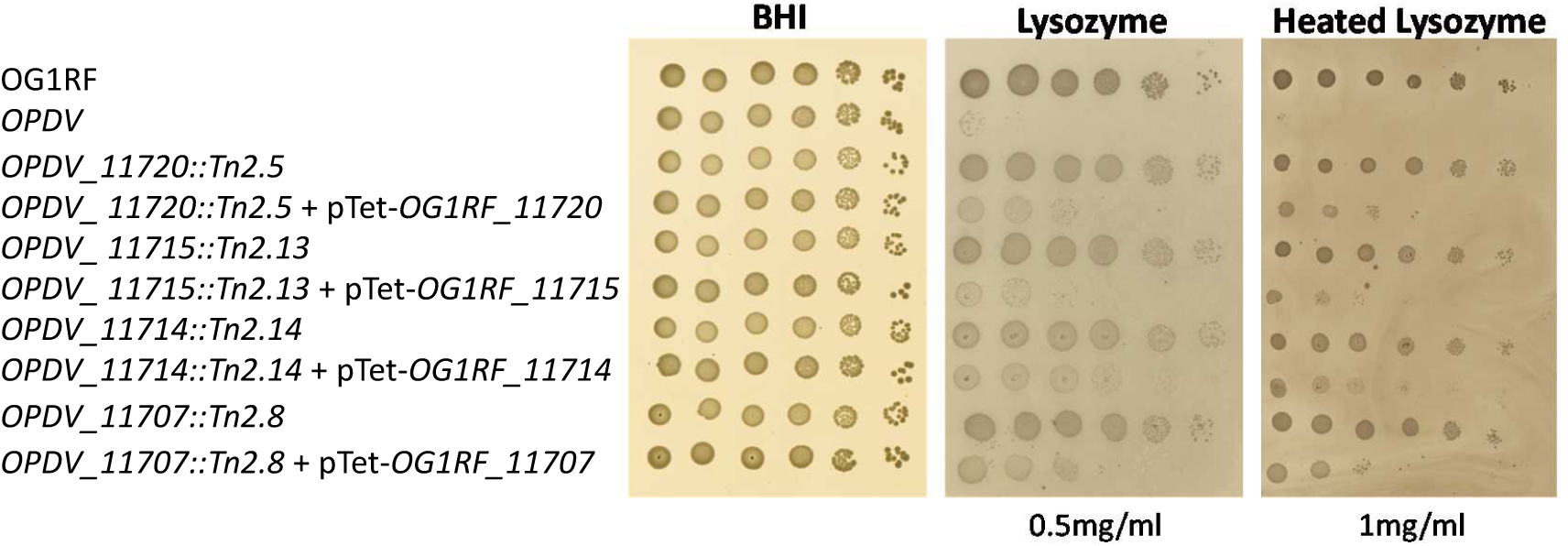
Growth defect of *E. faecalis* Tn mutants in the presence of antimicrobials targeting the cell envelope. Cell suspensions were prepared as described in S1 Fig and 1.5 µl of serial dilutions were spotted on BHI-agar plates containing 10 ng ml^−1^ anhydrotetracycline and various concentrations of lysozyme. Concentrations showing a clear difference in susceptibility are shown.

**Table 2.** Complementation of *epa* transposon mutants.

| Mutant | Complementation gene |  |  |  |
| --- | --- | --- | --- | --- |
|  | OG1RF_11720 | OG1RF_11715 | OG1RF_11714 | OG1RF_11707 |
| OPDV_11720::Tn2.9 | + | ND | ND | ND |
| OPDV_11720::Tn2.5 | + | - | - | - |
| OPDV_11715::Tn2.10 | ND | + | ND | ND |
| OPDV_11715::Tn2.13 | - | + | - | - |
| OPDV_11715::Tn2.16 | - | + | - | - |
| OPDV_11714::Tn2.14 | - | - | + | - |
| OPDV_11720::Tn2.12 | ND | ND | ND | + |
| OPDV_11720::Tn2.8 | ND | ND | - | + |
| OPDV_11720::Tn2.6 | ND | ND | ND | + |

<sup>a</sup>, Complementation was assessed on BHI-agar plates containing lysozyme at a concentration of 0.5 mg ml<sup>-1</sup>
+, complementation restoring lysozyme sensitivity; -, no impact on lysozyme MIC;
ND, not determined.

### Mutations in the *epa* variable region alter the negative surface charge of *E. faecalis* and are associated with minor changes in sugar composition

The impact of *epa* transposon insertions on the production of EPA was investigated. Polysaccharides were extracted from cultures at the end of exponential growth as previously described [25]. Similar amounts of EPA were extracted as assessed by neutral sugar assays and dry weight (between 20 and 30 mg l^−1^). Each purified EPA sample was run on a polyacrylamide gel and stained with the cationic dye alcian blue (Fig 3A). Whilst OG1RF and *OPDV* polysaccharide bands previously named PS1 and PS2 were present [8], these were not detected in mutant samples, possibly because the reduced negative charge no longer allowed EPA to migrate in the gel or no longer allowed these polymers to be stained by alcian blue. As expected, complementation restored the detection of EPA after staining by alcian blue (Fig 3A).

**Figure 3.**
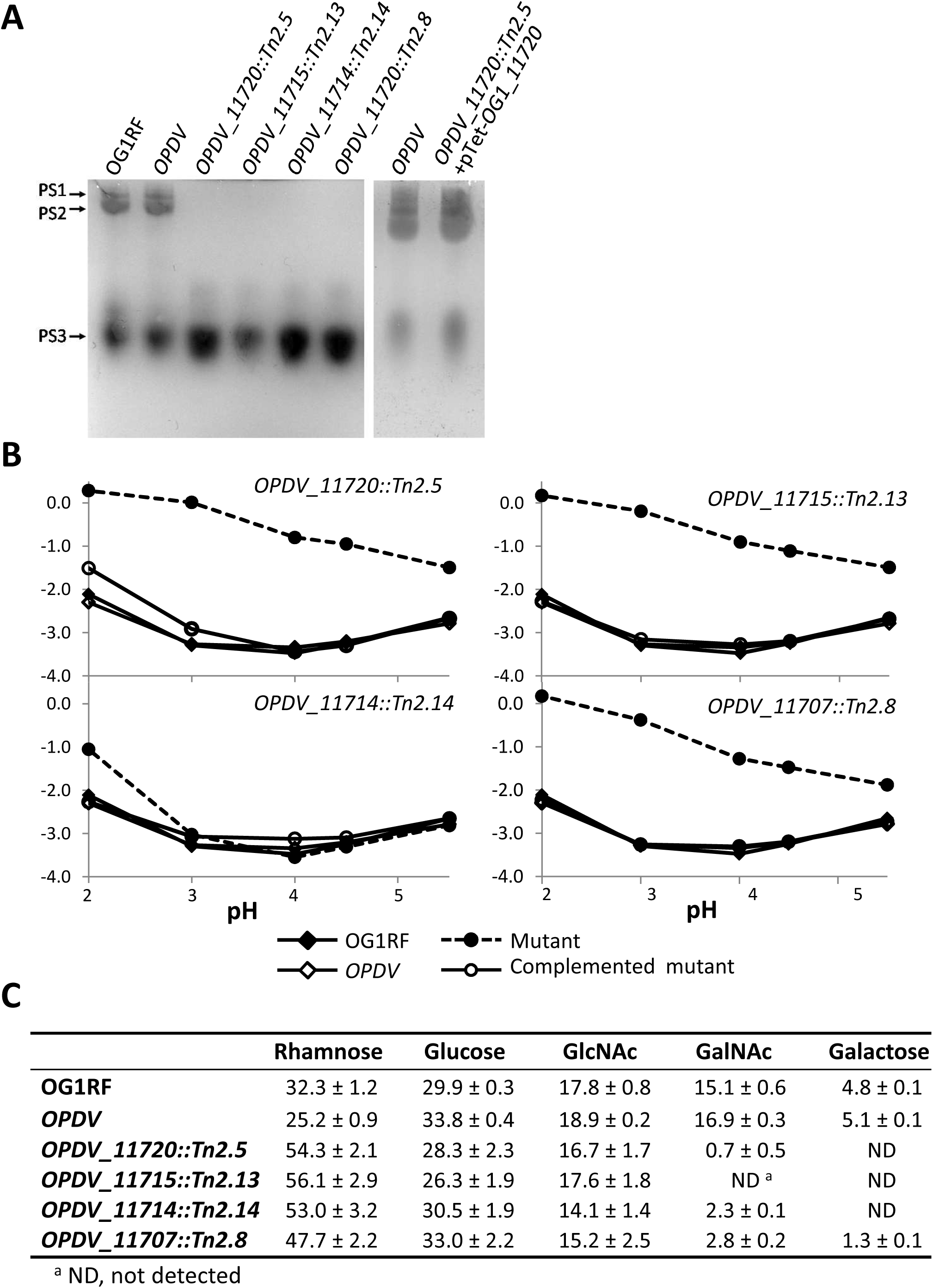
Analysis of purified EPA polysaccharides and their contribution to cell surface charge. **A.** Analysis of purified EPA by acrylamide gel electrophoresis. 40 µg of material was loaded on a 10 % (v/v) acrylamide-bisacrylamide (33:0.8) gel and stained with the cationic dye alcian blue. **B.** Electrophoretic mobility of *E. faecalis* OG1RF, *OPDV* and insertion mutants resistant to lysozyme. Representative mutants harbouring a transposon insertion in *OG1RF_11720* (*OPDV_11720::Tn2.5*), *OG1RF_11715* (*OPDV_11715::Tn2.13*), *OG1RF_11714* (*OPDV_11714::Tn2.14*) or *OG1RF_11707* (*OPDV_11707::Tn2.8*) were analysed. Wild-type OG1RF and parental *OPDV* strains were included as controls. **C.** Carbohydrate composition of purified EPA polysaccharides. The relative percentage corresponding to each monosaccharide was determined from three independent extractions.

We tested the impact of the transposon insertions on EPA charge by measuring the electrophoretic mobility of *E. faecalis* cells using micro-electrophoresis, which allows single particle tracking (Fig 3B) [26]. OG1RF cells displayed a negative electrophoretic mobility (migration towards the anode), even at low pH, indicating a negative surface charge. Despite the increased susceptibility of *OPDV* cells to lysozyme, the electrophoretic mobility measured with this strain was not significantly different from the mobility of OG1RF cells at all pHs tested (Fig 3B; S1 and S2 Table). Three of the four insertion mutants (*OPDV_11720::Tn2.5, OPDV_11715::Tn2.13 and OPDV_11720::Tn2.8*) displayed similar electrophoretic mobilities, very distinct from the parental *OPDV* strain. The negative surface charge of these mutants was significantly reduced as compared to that of the parental *OPDV* strain (\*\*\*\**P*<0.0001 for all pH conditions), the difference being most prominent at pH 3.0 (Fig 3B). As expected, differences between *OPDV, OPDV_11720::Tn2.5, OPDV_11715::Tn2.13* and *OPDV_11720::Tn2.8* cells were abolished when the mutations were complemented. Mutant *OPDV_11714::Tn2.14* only showed a difference with the parental *OPDV* strain at pH 2.0 (\*\*\*\**P*<0.0001). This difference was no longer detected when the mutation was complemented.

To confirm that the lack of detection of EPA on polyacrylamide gels was due to a loss of negatively charged groups rather than a lack of rhamnopolysaccharide production, we carried out carbohydrate composition analyses on purified EPA (Fig. 3C). As anticipated, EPA composition was very similar in OG1RF and *OPDV* strains; rhamnose, glucose, *N*-acetylglucosamine (GlcNac) and *N*-acetylgalactosamine (GalNAc) accounted for approximately 95% of the sugars identified (*ca.* 30%, 30%, 20% and 15%, respectively) and galactose was found in limited amounts (5%). The proportion of glucose and GlcNAc remained similar to parental *OPDV* levels in all *epa* mutants, whilst both GalNAc and galactose amounts decreased to an increase of rhamnose. Interestingly, these changes were different depending on the mutant considered. For example, EPA extracted from *OPDV_11718::Tn2.8* was the only mutant EPA that still contained some galactose. The relative proportion of rhamnose increased in all mutants. GalNAc content decreased dramatically in all mutant EPA and could not be detected in *OPDV_11715::Tn2.13*.

### NMR analyses reveal that the *epa* variable region contributes to minor modifications of the EPA polysaccharide

To gain further insight into the contribution of *epa* variable genes to the structure of EPA, we carried out NMR analyses on purified polysaccharides. The 1D proton NMR spectra of all polysaccharides were overall similar but mutations in the *epa* variable region were associated with modifications in the anomeric region (Fig 4A). Clear differences were also detected in the relative intensities of methyl protons in the mutant spectra. For all mutants, the intensity of *N*-acetyl signals (1.9-2.2 ppm) decreased whilst the intensity of methyl protons corresponding to rhamnose residues (1.2-1.6 ppm) increased, suggesting a lower content of hexosamine in mutant EPAs and a relative increase of rhamnose. This result is in agreement with the carbohydrate analyses following acid hydrolysis (Fig 3C and S3 Fig). ^1^H-^13^C HSQC spectra revealed that EPA has a very complex structure, as evidenced by the detection of over 30 signals in the anomeric region (Fig 4B). The comparison of 2D spectra corresponding to *OPDV* and mutant polysaccharides revealed that each *epa* mutation only led to a limited number of changes including changes in the signal intensity, signal shifts and disappearance (S4 Fig). The number and the nature of the signals affected in the *epa* mutants were different depending on the mutation considered. Altogether, NMR and sugar analyses supported the idea that the *epa* variable genes are involved in limited modifications of the EPA rhamnopolysaccharide previously described as “decorations”.

**Figure 4.**
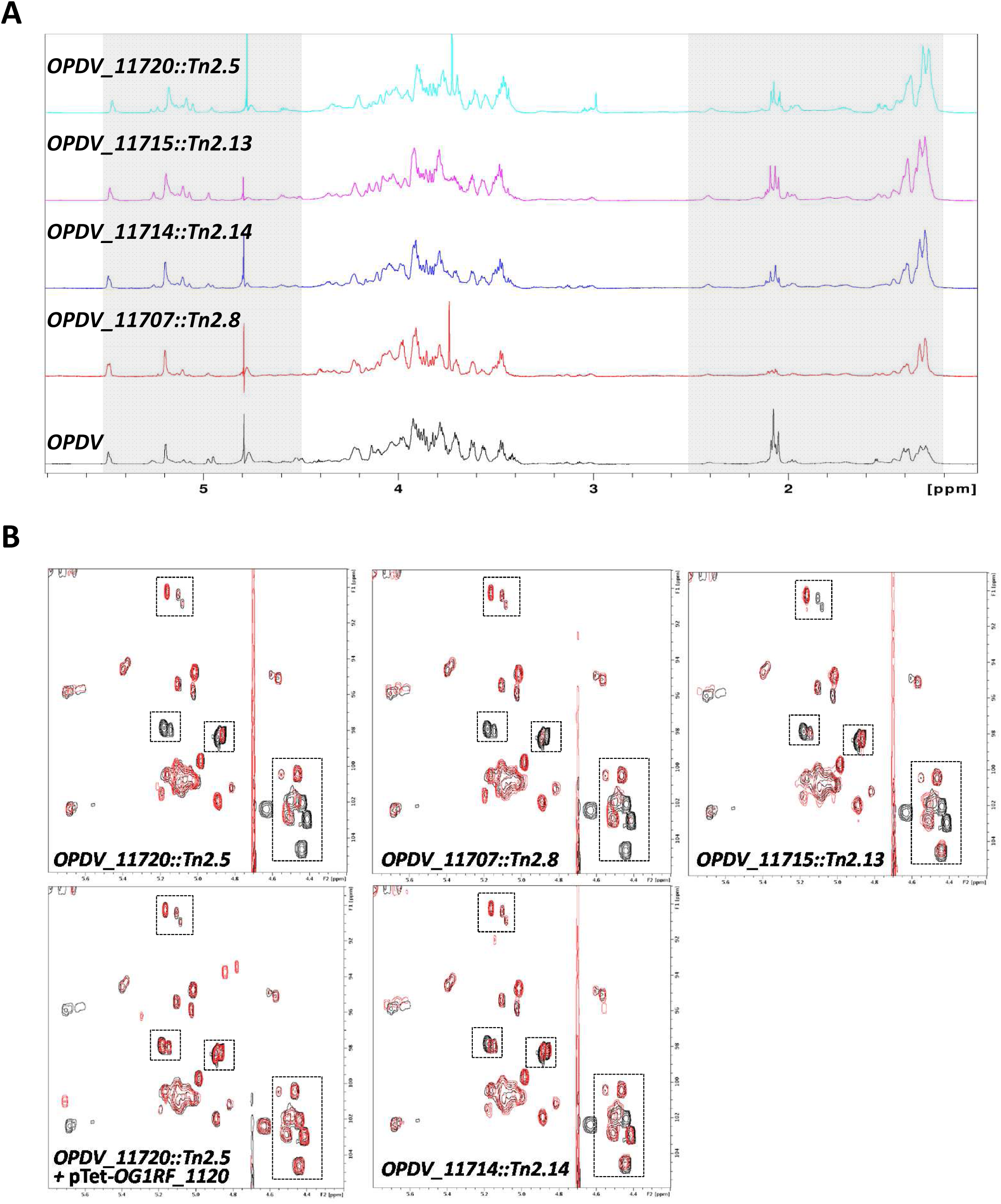
Structural analysis of purified EPA polysaccharides. **A.** 1D proton spectra of the EPA polysaccharides extracted from strains *OPDV, OPDV_11720::Tn2.5, OPDV_11715::Tn2.13, OPDV_11714::Tn2.14* or *OPDV_11707::Tn2.8*. The grey boxes indicate anomeric (4.5-5.5 ppm) and methyl protons (1.2-2.5 ppm). **B.** 2D ^1^H-^13^C HSQC spectra of EPA polysaccharides. The region corresponding to anomeric protons (4.2-5.5 ppm) and anomeric carbons (90-105 ppm) is shown. *OPDV* signals are in black, mutant signals in red. Boxes show signals with a lower intensity or a shift in the mutant EPA samples. Close-up views of the boxed regions are shown in S4 Fig.

### Epa decoration determines susceptibility to antimicrobials targeting the cell envelope

All *epa* transposon insertions were combined with four other mutations present in the in *OPDV* strain (*oatA, pgdA, dltA* and *sigV*), leaving the possibility of epistatic interactions between these mutations. To avoid this potential issue, we built in-frame *epa* deletions in the OG1RF genetic background (S5 Fig) before testing the impact of EPA decorations on resistance to antimicrobials targeting the cell envelope (Fig 5). All *epa* mutants were more sensitive to SDS than OG1RF, mutants Δ*11714* and Δ*11707* being less sensitive than the two others. Interestingly, mutant Δ*11714* (Δ*epaX-like*) was the only one that did not display increased susceptibility to sodium cholate, a primary bile salt. Mutant Δ*11720* was the only one with an increased susceptibility to both polymyxin B and nisin, two cationic peptides targeting the cell envelope. Mutants Δ*11707* and Δ*11715* were more susceptible to polymyxin B than the wild-type strain, but barely more susceptible to nisin. Finally, deletion of *OG1RF_11714* had no detectable impact on resistance to either of the CAMPs tested. Taken together, these results indicated that genes in the *epa* variable region are required for resistance to antimicrobials targeting the cell envelope.

**Figure 5.**
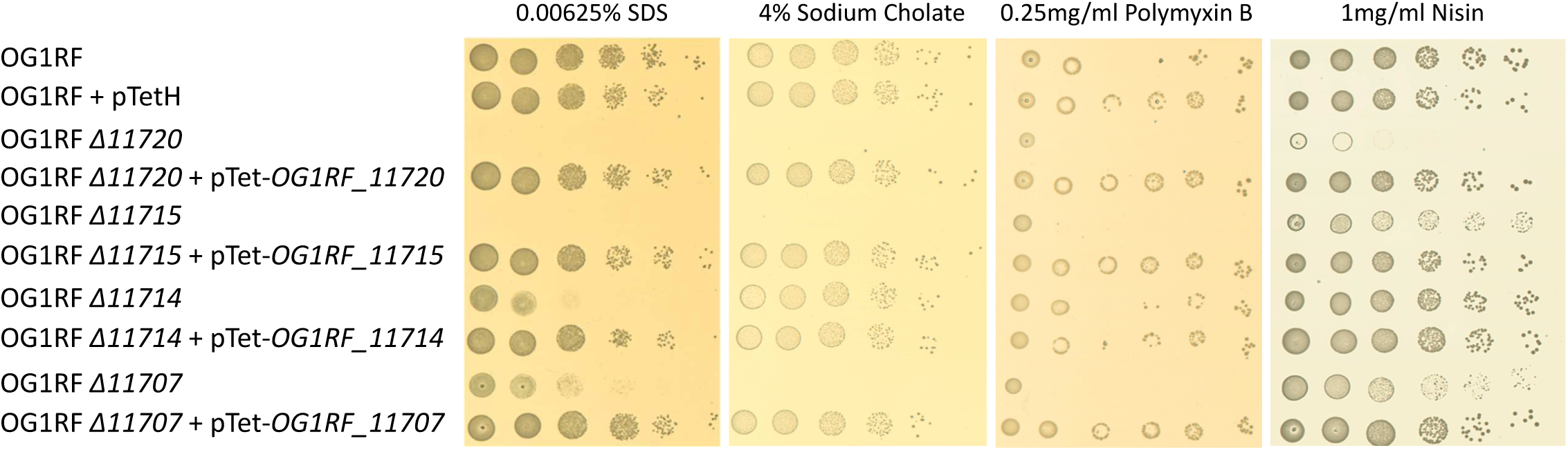
Growth defect of *E. faecalis* OG1RF in frame *epa* mutants in the presence of antimicrobials targeting the cell envelope. Cell suspensions were prepared as described in S1 Fig and 1.5 µl of serial dilutions were spotted on BHI-agar plates containing 10 ng ml^−1^ anhydrotetracycline and SDS, sodium cholate, polymyxin B or nisin. For each antimicrobial, one concentration showing a difference in susceptibility is shown.

### The decoration of the EPA rhamnopolysaccharide is essential for virulence and underpins phagocyte evasion

The impact of *epa* mutations on *E. faecalis* virulence was tested in the zebrafish experimental model of infection (Fig 6). Cell suspensions corresponding to approximately 1,200 CFUs were injected in the bloodstream of LWT embryos 30 h post fertilization (hpf) and the survival of larvae was monitored for 90 h post infection (hpi). As a preliminary experiment, we analysed the virulence of the *OPDV* transposon mutants (S6 Fig). Each *epa* transposon mutant had a significantly reduced virulence as compared to the wild-type OG1RF strain. Even though the combined deletions in *oatA, pgdA, dltA* and *sigV* did not impair the virulence of *E. faecalis* in the zebrafish model of infection (S7 Fig), we could not exclude the possibility of an epistatic relation between the *OPDV* and *epa* mutations. We therefore repeated the zebrafish infections using the in-frame *epa* deletion mutants in the wild-type OG1RF genetic background (S5 Fig). All *epa* mutants showed a significant decrease in virulence as compared to the wild-type OG1RF (Fig 6), killing only between 0-10% of the larvae as opposed to the 40 to 55% of killing following injection of the wild-type strain; Fig 6A-D). As expected, the complementation of each *epa* deletion fully restored the virulence. Although the *epa* deletion mutants (except the Δ*epaX-like* strain) present a slight defect in their growth rate, it is unlikely that this accounts for the lack of virulence; all complemented strains (and the wild-type OG1RF harbouring an empty complementation vector) also present a growth defect (S8 Fig) and yet kill zebrafish larvae as well as the wild-type strain (Fig 6).

**Figure 6.**
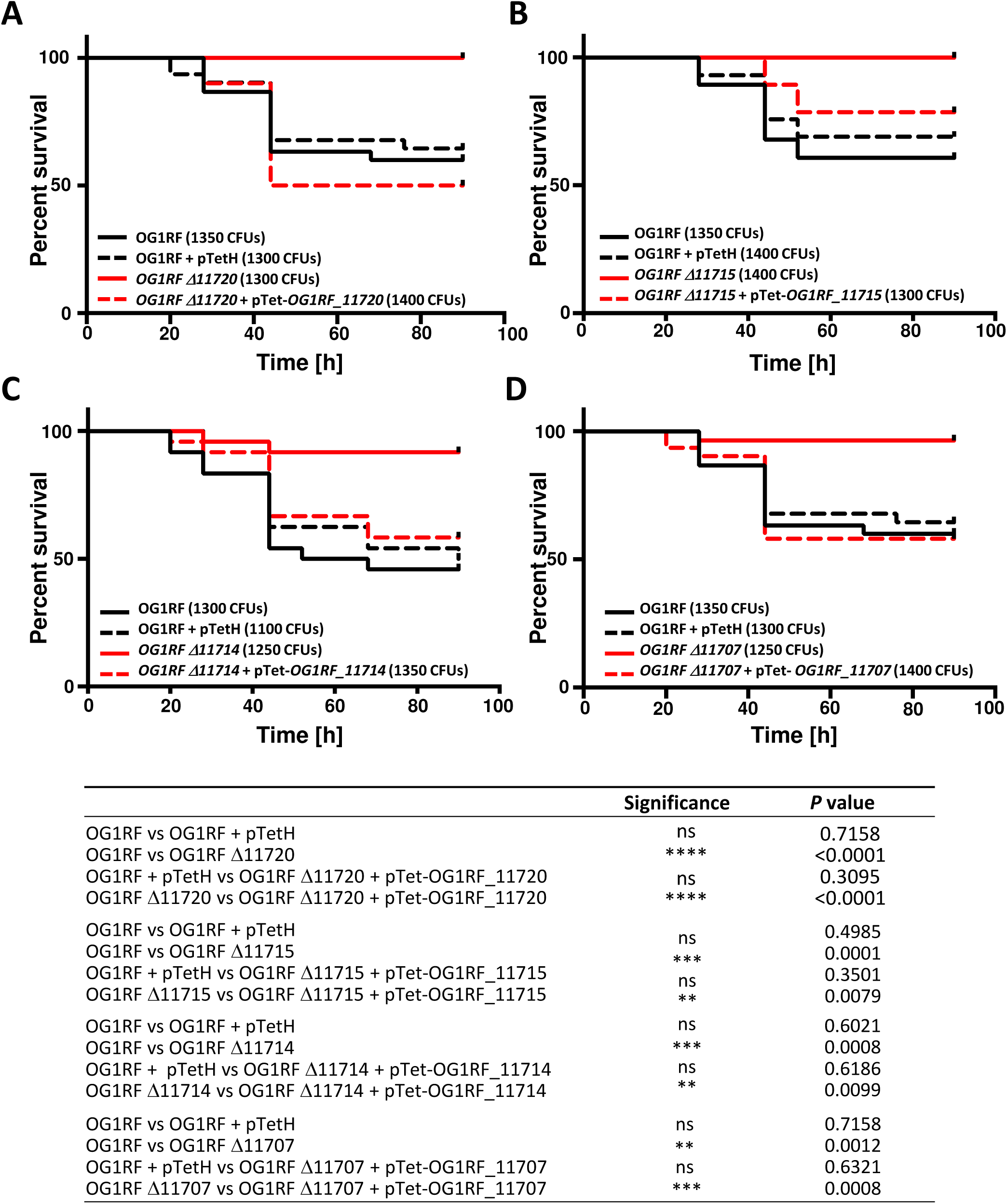
Virulence of *epa* mutants and complemented strains in the zebrafish model of infection. Survival of zebrafish larvae (n>20) following infection with *E. faecalis* OG1RF (WT) and *epa* deletion mutants was monitored over 90 h post infection. **A.** Mutant Δ*11720*. **B.** Mutant Δ*11715*. **C.** Mutant Δ*11714*. **D**. Mutant Δ*11707*. **E.** Statistical significance determined by Log-rank test; ns, *P*>0.05; \*\**P*<0.01; *** *P*<0.001; **** *P*<0.0001. All injections presented in Fig 4A and 4D were carried out on the same day. The same data corresponding to the OG1RF strain are therefore shown for the 2 experiments.

The production of EPA has been associated with an increased resistance to phagocytosis, which represents a key step during pathogenesis [8, 27]. We therefore quantified phagocytosis in zebrafish larvae infected with the wild-type OG1RF and one representative *epa* mutant (*OG1RF_*Δ*11714, epaX-like*) expressing the red fluorescent protein mCherry (Fig 7). Confocal microscopy images were used to measure bacterial uptake by phagocytes labelled with anti L-plastin antibodies coupled to Alexa-488, a green fluorophore, as previously described [28]. The ratio between red fluorescence inside to red fluorescence outside phagocytes was significantly higher for the *epaX-like* mutant (*OG1RF_*Δ*11714*) than for the wild-type strain (\*\*\**P*=0.0006) or the complemented strain (OG1RF Δ*11714* + pTetH-*OG1RF_11714*; \*\**P* = 0.0049). As expected, no difference in phagocytosis was observed between the wild-type and complemented strains (ns, *P*>0.05) (Fig 7A). Representative pictures shown in Fig 7B-D clearly indicate that unlike the wild-type strain, the *epaX-like* mutant was no longer able to evade phagocytes.

**Fig 7.**
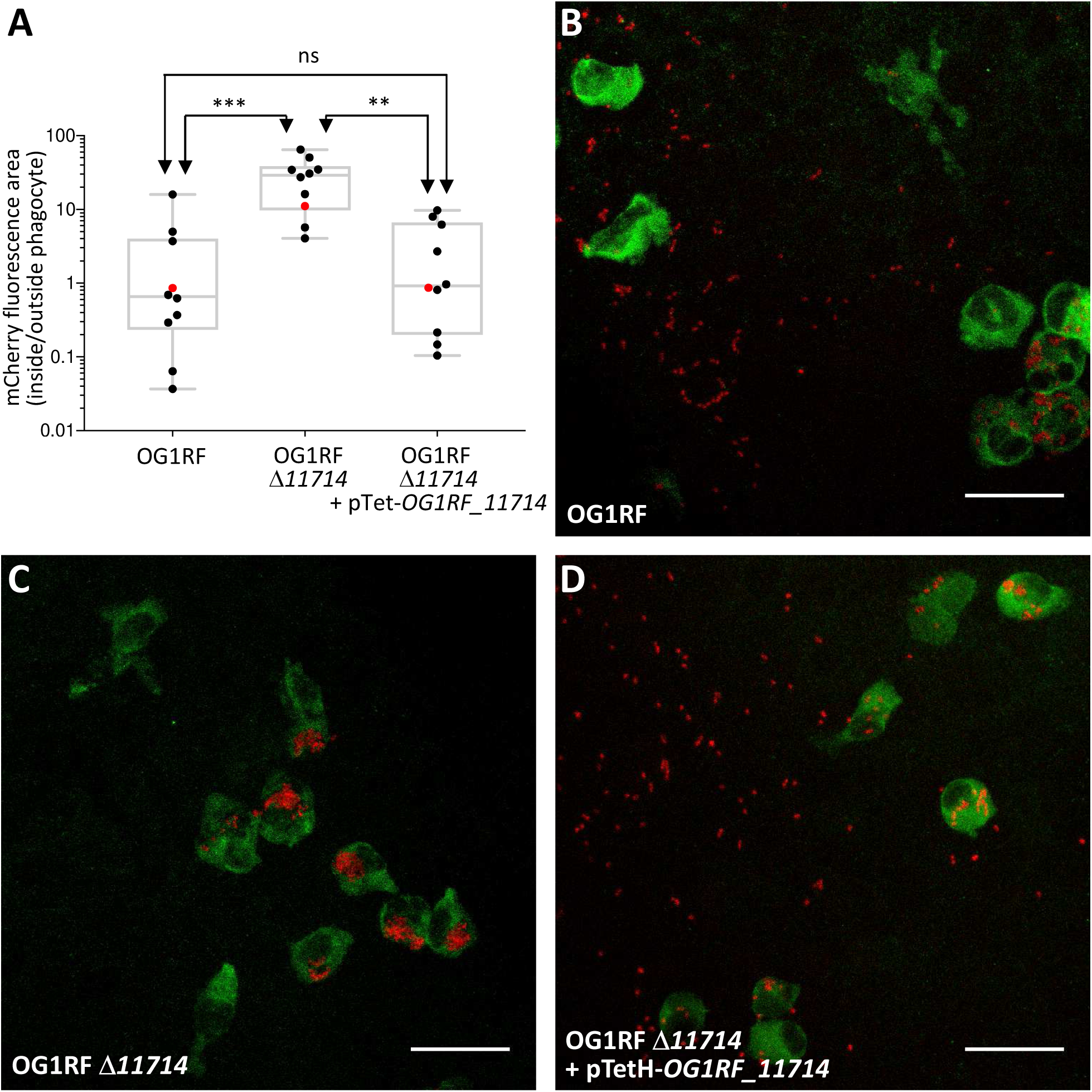
*epaX* mutant cells are more prone to phagocytosis than wild-type and complemented cells. **A.** Quantification of *E*. *faecalis* uptake by zebrafish phagocytes. Embryos were infected with 1,600 CFUs of *E*. *faecalis* cells constitutively producing mCherry and fixed in 4 % paraformaldehyde 1.5 h post infection. Phagocytes were immunolabelled using rabbit anti L-plastin antibodies and detected with goat anti-rabbit antibodies conjugated to Alexafluor 488. The infected and immunolabelled embryos were imaged using scanning confocal microscope. The ratio of mCherry fluorescence signal area associated with phagocytosed and free bacteria was measured using the Fish Analysis Fiji plugin. The uptake of mutant OG1RF Δ*11714* (Δ*epaX*) was significantly higher when compared to the wild-type (OG1RF; \*\*\**P* = 0.0006) and complemented strain (OG1RF Δ*11714* + pTetH-*OG1RF_11714*; \*\**P* = 0.0049). No difference in phagocytosis was observed between the wild-type and complemented strains (ns, *P*>0.05). Representative images showing *E. faecalis* uptake in zebrafish embryos are shown. Each picture corresponds to the quantification result indicated with a red dot in **A**, following infection with OG1RF (**B**), OG1RF Δ*11714* (Δ*epaX*) (**C**) and the complemented strain (**D**). Phagocytes labeled with L-plastin appear in green, mCherry labelled bacteria in red. Scale bar is 25 μm.

Altogether, our data therefore indicate that decoration of the EPA polysaccharide is essential for *E. faecalis* pathogenesis.

## Discussion

Previous studies revealed that *E. faecalis* resistance to lysozyme is unusually complex and results from several mechanisms acting synergistically. These include peptidoglycan *O*-acetylation and de-*N*-acetylation, D-alanylation of teichoic acids and transcriptional control by the extracytoplasmic sigma factor SigV. Despite a 100-fold decrease as compared to the wild-type strain, the residual resistance of the quadruple *OPDV* mutant is still relatively high (MIC=0.5 mg ml^−1^) as compared to other Firmicutes. This result contrasts with other bacteria in which a limited number of genes play a key role in resistance. For example, the combined deletions of *oatA* and *dltA* in *S. aureus* led to a decrease of at least 2,000-fold in resistance as compared to parental strain [13]. Deletion of *pgdA* alone in *L. monocytogenes* is associated with a 32-fold decrease resistance.

Using random transposon mutagenesis, we showed that several genes located downstream the conserved *epaA-epaR* region modulate susceptibility to lysozyme. It was proposed that *epaA-epaR* encode a core synthetic machinery whilst downstream genes contribute to the decoration of EPA rhamnopolysaccharide [22, 24]. In agreement with previous studies, three distinct polysaccharide bands (named PS1, PS2 and PS3) were detected on polyacrylamide gels ([8] and Fig 3A). The two upper bands simultaneously disappeared in all the transposon mutants, suggesting that both are structurally related to EPA. The nature of the third band is unknown and could be either a metabolic intermediate of EPA or an unrelated polymer. The lack of detection of PS1 and PS2 in EPA polysaccharides from mutants suggested that either their charge did not allow them to enter the gel and/or that they were no longer stained by the cationic dye alcian blue. A similar result was previously described for mutants harbouring deletions in *OG1RF_11715* (*epaOX*) [23] and the homolog of *OG1RF_11714* in V583 (*epaX*) [24]. By comparing the electrophoretic mobility of the transposon mutants to that of the parental *OPDV* strain, we confirmed that the *epa* genes downstream of the *epaA-epaR* locus contribute to the negative charge of the EPA polysaccharide. This negative charge could at least in part be due to the presence of phosphate in the polymer. Interestingly, the OG1RF and *OPDV* strains displayed similar electrophoretic mobilities. Previous studies also showed that alanylation of teichoic and/or lipoteichoic acids in *L. lactis* had no detectable impact on the electrophoretic mobility of this organism [29]. The *dlt* operon has been shown to modify lipoteichoic acids [17]. Since these polymers are embedded inside the cell wall, it is likely that their modification does not lead to a change in the bacterial surface charge. Further experiments are required to test whether alanylation of cell wall polymers only has a moderate impact on the charge of the cell wall or if such modifications can simply not be detected by measuring electrophoretic mobility. Three of the mutants identified in this work carry a transposon in genes encoding putative glycosyl transferases. Despite the low amino acid identity between the glycosyl transferase sequences (19-27% depending on the comparison), these proteins have very similar predicted secondary structures, with two transmembrane domains and both the N- and C-termini exposed at the cell surface. Tertiary structure predictions suggest that all 3 proteins have a very similar fold and are GalNAc transferases. These predictions are in agreement with our NMR and sugar analyses indicating that EPA polysaccharides from all glycosyl transferase mutants (*OPDV_11720, OPDV_11715* and *OPDV_11714*) contain less GalNAc and less intense *N*-acetyl proton signals. In addition to a reduced amount of GalNAc, EPA polymers from the glycosyl transferase mutants also contained a reduced amount of galactose. Further analyses are required to explore the catalytic activity of these glycosyl hydrolases. It remains unclear whether they can use distinct sugars as a substrate or if the addition of GalNAc is required for the activity of other glycosyl transferases adding Gal residues. Since none of the heterologous complementations of the transposon mutants were able to restore the parental phenotype, we anticipate that the 3 glycosyl transferases identified play distinct roles. This idea is supported by several independent observations: (i) mutants *OPDV_11720::Tn2.5, OPDV_11715::Tn2.13* and *OPDV_11714::Tn2.14* present distinct alterations in their EPA carbohydrate compositions and the deletion mutants present differences in their antimicrobial susceptibility profiles; (ii) 2D-NMR spectra indicate that each mutation is associated with distinct modifications of the signals in the anomeric region; (iii) the *OPDV_11714::Tn2.14* (*epaX-like*) mutant behaved differently from the other *epa* mutants studied since it displayed a less pronounced defect in surface charge. Altogether, our results suggest that glycosyltransferases in the *epa* variable region fulfil distinct roles. The complexity of EPA structure precludes any conclusion about the specific role of individual *epa* genes. However, based on the 2D NMR spectra, it is tempting to assume that *OG1RF_11720*, the first gene of the *epa* variable region encodes a glycosyl transferase that is adding the entire EPA decorations, whilst EpaOX and EpaX contribute to the transfer of smaller decorations.

*epa* deletion mutants displayed distinct resistance to antimicrobials targeting the cell envelope. Deletion of *OG1RF_11720,* which had the most pronounced impact on EPA structure led to the most pronounced increased susceptibility to antimicrobials. Deletion of *OG1RF_11714* (*epaX*), which had the least pronounced impact on EPA structure only led to an increased susceptibility to SDS. Since *epa* mutations could confer resistance to both negatively charged antimicrobials and CAMPs, these results suggested that the charge of EPA does not entirely account for the resistance to these compounds. We therefore speculate that EPA decorations are critical to maintain cell integrity, as previously suggested [23, 24]. A contribution of *epa* decoration genes in biofilm formation [30], resistance to antimicrobials [23] and colonisation [24] was previously suggested, but no information was available on a potential role in the context of pathogeny. *E. faecalis* pathogenesis in the zebrafish model of infection involves two critical steps: phagocyte evasion and tissue damage caused by the metalloprotease GelE [27]. Since *oatA, pgdA, dltA* and *sigV* are unlikely to contribute to these processes, it was expected that their simultaneous deletion would have a very limited impact on virulence. By contrast, EPA has been reported to mediate resistance to phagocytic killing [8] and plays a critical role in virulence both in experimental mouse and zebrafish infections [27, 31]. We therefore hypothesized that *epa* mutations altering the decoration of EPA would impair virulence. In agreement with this hypothesis, all *epa* transposon mutants (in the *OPDV* background) and in-frame deletions in the wild-type OG1RF background were avirulent in the zebrafish model of infection. Further investigations revealed that the *epaX* mutation leads to a significant increase in *E. faecalis* uptake by phagocytes, suggesting that the decoration of EPA mediates immune evasion and underpins virulence.

Collectively, the results provide a paradigm shift in our understanding of *E. faecalis* pathogenesis, revealing that the modifications of EPA, rather than EPA backbone itself, underpin phagocyte evasion, an essential step during host infection. Whether *epa* is directly recognized by the host immune system or is shielding other surface components remains an open question.

## Materials and methods

### Ethics statement

Animal work was carried out according to guidelines and legislation set out in UK law in the Animals (Scientific Procedures) Act 1986 under Project License P1A417A5E. Ethical approval was granted by the University of Sheffield Local Ethical Review Panel.

### Bacterial strains, plasmids and growth conditions

Bacterial strains, plasmids and oligonucleotides used in this study are described in S3 Table. All strains were routinely grown at 37°C in Brain Heart Infusion (BHI) broth or BHI-agar 1.5 % (w/v) plates unless otherwise stated. For *E. coli*, erythromycin was added at a final concentration of 200 µg ml^−1^ to select pTetH derivatives. When necessary, *E. faecalis* was grown in the presence of 10 µg ml^−1^ chloramphenicol, 128 µg ml^−1^ gentamicin or 30 µg ml^−1^ erythromycin. For complementation experiments, anhydrotetracycline was used at a final concentration of 10 ng ml^−1^ to induce gene expression.

### Construction of the *OPDV* strain

The work describing the contribution of *oatA, pgdA, dltA* and *sigV* to lysozyme resistance was carried out using *E. faecalis* JH2-2 as a genetic background [16, 19]. Since this laboratory strain is avirulent in the zebrafish model of infection, we decided to use OG1RF as a parental strain [32]. The quadruple OG1RF mutant harbouring deletions in *oatA, pgdA, dltA* and *sigV* was built using existing plasmids (S3 Table) to create in-frame deletions in the following order: *oatA, pgdA, dltA* and *sigV*.

### Transposon mutagenesis

A *Mariner*-based transposon mutagenesis system previously described was used [21]. Plasmid pZXL5 was introduced in *E. faecalis OPDV* by electroporation and transformants were selected at 28°C on plates containing chloramphenicol and gentamicin. Cells harbouring pZXL5 were grown to mid-exponential phase at 28°C and transposition was induced by addition of nisin (25 ng ml^−1^). The culture was then transferred to 42°C overnight to counter-select the replication of the plasmid. The library was then amplified by growing the cells at 42°C in the presence of gentamicin.

### Transposon mutagenesis

A *Mariner*-based transposon mutagenesis system previously described was used [21]. Plasmid pZXL5 was introduced in *E. faecalis OPDV* by electroporation and transformants were selected at 28°C on plates containing chloramphenicol and gentamicin. Cells harbouring pZXL5 were grown to mid-exponential phase at 28°C and transposition was induced by addition of nisin (25 ng ml^−1^). The culture was then transferred to 42°C overnight to counter-select the replication of the plasmid. The library was then amplified by growing the cells at 42°C in the presence of gentamicin.

### Isolation of transposon mutants resistant to lysozyme

Serial dilutions of the transposon library were plated on BHI agar plates containing 1, 2 or 4 times the lysozyme MIC for the *OPDV* strain (0.5 mg ml^−1^) and gentamicin. After 24 to 48 h at 42°C, individual colonies growing at the highest concentration (2 mg ml^−1^) were chosen for further characterisation.

### Mapping transposition sites

Transposon insertion sites were mapped by reverse PCR using two divergent primers (Mar_up and Mar_dn) on the transposon (S9A Fig). Chromosomal DNA was extracted using the Promega Wizard kit and digested by SspI in a final volume of 30 µl at a concentration of 4 ng µl^−1^ (S9B Fig). Digestion products were further diluted to 1 ng µl^−1^ and self-ligated at 16°C for 16 h after addition of 100 U of T4 DNA ligase (NEB) (S9C Fig). Three microliters of the ligation product were used as a template for PCR amplification using oligonucleotides Mar_up and Mar_dn (S9D Fig). PCR products were gel extracted and sequenced using oligonucleotide T7 (S9E Fig). The insertion site was defined as the first nucleotide of the *E. faecalis* OG1RF genome immediately downstream of the inverted repeat sequence flanking the transposon.

### Construction of complementation plasmids

DNA fragments encoding OG1RF_11720, OG1RF_11715, OG1RF_11714 and OG1RF_11707 were amplified by PCR using the oligonucleotides described in S3 Table. PCR products were digested by NcoI and BamHI and cloned into pTetH, a pAT18 derivative allowing anhydrotetracycline-inducible expression (S. Mesnage, unpublished). Each open reading frame was fused to a C-terminal 6-Histidine tag.

### Antimicrobial assays

Colonies from a BHI agar plate were resuspended in PBS and diluted to an OD of 1 at 600 nm. Ten-fold dilutions were prepared in PBS and 1.5 µl of each cell suspension were spotted on BHI agar containing antimicrobials at various concentrations. MICs of lysozyme were defined as the concentration of antimicrobial inhibiting the growth of 1.5 µl of a cell suspension corresponding to a 1000-fold dilution of the cell suspension at OD of 1 at 600 nm. For complementation experiments, anhydrotetracycline was added at a final concentration of 10 ng ml^−1^. At least two biological replicates were carried out for each susceptibility assay.

### Measurement of electrophoretic mobility

An overnight culture was diluted 1000-fold in 25 ml of BHI broth and grown for 17 h at 37°C in static conditions. Anhydrotetracycline (100 ng ml^−1^) was added to all cultures to induce gene expression for complementation and exclude the possibility that this chemical could account for differences between strains. Cells were harvested by centrifugation (5 min, 8,000 x *g* at room temperature), washed twice in 25 ml of 1.5 mM NaCl and resuspended at a concentration of 3×10^7^ CFU ml^−1^ in 1.5 mM NaCl at various pHs. The electrophoretic mobility was measured in an electric field of 8 V cm^−1^ using a laser zetaphoremeter (CAD Instrumentation, Les Essarts le Roy, France). For each measurement, results were based on the analysis of 200 individual particles. The results presented in Fig 3, S1 and S2 Tables are the combined results of three independent experiments (biological replicates).

### NMR

NMR experiments were conducted on a Bruker DRX-600 (plus cryoprobe) spectrometer at 25°C. EPA polysaccharides were freeze-dried and resuspended in D_2_O. Spectra were processed and analysed using TOPSPIN (version 2.1). Trimethylsilylpropanoic acid was used as a reference.

### Carbohydrate extraction and analyses

EPA was extracted as previously described from standing cultures in BHI at the end of exponential growth (OD_600nm_=0.8) [25]. The method previously described to analyse pneumococcal polysaccharides and conjugates was followed, with the exception of the first acid hydrolysis step [33] Briefly, purified EPA polysaccharides were hydrolyzed in 4 N trifluoroacetic acid for 4 h at 100°C. Hydrolysis products were analysed by high-performance anion-exchange chromatography (HPAEC) coupled to pulsed-amperometric detection (PAD) using a Dionex DX 500 BioLC system (ThermoFisher). Monosaccharides were separated on a Carbopac PA10 (4 mm × 250 mm) analytical column (Thermofisher Scientific) at a flow rate of 1 ml min^−1^. Solvent A was 18 mM NaOH, solvent B was 100 mM NaOH, and solvent C was 100 mM NaOH containing 1 M sodium acetate. NaOH and NaAc gradients were used simultaneously to elute the carbohydrates by mixing the three eluents. The gradients used were as follows: after 15 min of isocratic elution in buffer A, a 3 min gradient to 100 % of buffer B was applied. A second gradient was applied between 18 and 35 min using buffer C to reach 300 mM sodium acetate. The column was re-equilibrated in 18 mM NaOH for 20 min after every run. The following pulse potentials and durations were used: *E*1 = 0.1V, *t*1 = 400 ms; *E*2 = −2V, *t*2 = 20 ms; *E*3 = 0.6V, *t*3 = 10 ms; *E*4 = −0.1V, *t*4 = 70 ms. Data were collected and analysed on computers equipped with the Dionex PeakNet software. Carbohydrate analyses were made in triplicate using three independent EPA extractions from 3 distinct colonies.

### Construction of pGhost derivatives for allele replacement

All plasmids for allele replacement were constructed with the same strategy. Two homology regions were amplified: the 5’ homology region (referred to as H1) was amplified with oligonucleotides H11 (sense) and H12 (antisense). The 3’ homology region (referred to as H2) was amplified with oligonucleotides H21 (sense) and H22 (antisense). Both PCR products were purified, mixed in an equimolar ratio and fused by overlap extension using oligonucleotides H11 and H22 [34]. The assembled PCR fragment flanked by two restriction sites was digested and cloned into pGhost9 [35] similarly digested. Oligonucleotide sequences and restriction sites used for cloning are described in S3 Table.

### Construction of *E. faecalis* OG1RF in-frame *epa* mutants

Isogenic derivatives of *E. faecalis* OG1RF were constructed by allele exchange using the procedure previously described [36]. Briefly, pGhost9 derivatives were electroporated into OG1RF and transformants were selected at a permissive temperature (28°C) on BHI plates with erythromycin. To induce single crossover recombination, transformants were grown at a non-permissive temperature (42°C) in the presence of erythromycin. The second recombination event leading to plasmid excision was obtained after 5 serial subcultures at 28°C without erythromycin. The last overnight subculture was plated at 42°C without erythromycin. A clone harboring a double crossover mutation was identified by PCR (S5 Fig) and further confirmed by sequencing of the recombined region.

### Zebrafish strains and maintenance

London wild type (LWT) zebrafish were provided by the aquarium facility at the University of Sheffield. Embryos were maintained in E3 medium at 28°C according to standard procedures previously described [37].

### Microinjections of *E. faecalis* in zebrafish embryos

Cells were grown to mid-exponential phase (OD_600nm_∼0.3) and harvested by centrifugation (5,000 × *g* for 10 min at room temperature). Bacteria were resuspended in filtered phosphate buffer saline (150 mM Na_2_HPO_4_, 20 mM KH_2_PO_4_, 150 mM NaCl [pH 7.5], PBS) and transferred to microcapillary pipettes. Embryos at 30 h post fertilization (hpf) were anaesthetized, dechorionated, embedded in 3 % (w/v) methylcellulose and injected individually with 2 nl of a cell suspension corresponding to *ca.* 1,000 cells as previously described [27]. The number of cells injected was checked before and after each series of injections with a given strain. Zebrafish embryos were monitored at regular intervals until 90 h post infection (hpi). At least 20 embryos per group were used.

### Imaging of infected larvae by confocal microscopy and quantification of uptake by phagocytes

Immuno-labelled embryos were immersed in 0.8 % (w/v) low melting point agarose in E3 medium and mounted flat on FluoroDish™ (World Precision Instruments Inc.). Images were collected using a DMi8 confocal microscope (Leica). Image acquisition was performed with the Volocity software and the images were processed with ImageJ 1.49v software. Bacterial phagocytosis was quantified using an ImageJ custom script called Fish Analysis, which can be obtained from http://sites.imagej.net/Willemsejj/ or via ImageJ updater. All bacteria were identified based on their fluorescence (mCherry, Channel 2). Subsequently, the fluorescence intensity of the phagocytes (Alexa 488, Channel 1) surrounding the phagocytosed bacteria was measured. The phagocytosed bacteria had high fluorescence intensity of Channel 2 and low fluorescence intensity of Channel 1. The area of phagocytosed bacteria was compared with the area of non-phagocytosed bacteria and their ratio was calculated.

### Statistical analyses

Statistical analyses were performed using GraphPad Prism version 7.03. Comparisons between survival curves were made using the log rank (Mantel-Cox) test. Electrophoretic mobilities were compared using two-way ANOVA Comparison of uptake by zebrafish macrophages was carried out using an unpaired non-parametric Dunn’s multiple comparison test.

## Supporting information

## Acknowledgments

The authors would like to thank Abdellah Benachour (University of Caen) for providing the plasmids used to construct the *OPDV* strain and the Bateson Centre aquaria staff for their assistance with zebrafish husbandry. The authors thank Dr Philip Elks for access to the confocal microscope and Paul Martin (University of Bristol) for the kind gift of anti L-plastin antibodies.

## Supporting information Legend

**S1 Figure. Lysozyme MICs for Firmicutes.** A cell suspension in phosphate saline buffer was adjusted to an OD at 600 nm of 1 and 1.5 µl of serial dilutions were spotted on BHI-agar plates containing various concentrations of lysozyme. ND, undiluted cell suspension; 10^−1^, 10-fold dilution; 10^−2^, 100-fold dilution; 10^−3^, 1000-fold dilution; 10^−4^, 10000-fold dilution; 10^−5^, 100000-fold dilution.

**S2 Figure. Western blot analysis of complementation strains.** Cultures were grown in BHI to an OD at 600 nm of 0.5 and expression of the *epa* genes was induced by addition of anhydrotetracycline (10 ng ml^−1^). After 2 h, cells were harvested and mechanically broken in the presence of glass beads. Crude extracts (20µg) were loaded on SDS-PAGE, transferred onto a nitrocellulose membrane and probed with a polyclonal serum against the polyhistidine tag. Bands of the expected molecular weights were detected (OG1RF_11707, 36.7 kDa; OG1RF_11714, 38.9 kDa; OG1RF_11715, 38.4 kDa; OG1RF_11720, 30.8 kDa).

**S3 Figure. HPLC analysis of TFA-hydrolysed EPA polysaccharides.** Following gel filtration, fractions containing neutral sugars were pooled and freeze-dried. EPA was hydrolysed in the presence of 4 N TFA at 100°C for 4 h. Monosaccharides were separated on a carbopac PA10 column by high performance anion exchange chromatography coupled to pulsed-amperometric detection. Representative chromatograms are shown for monosaccharide standards and each transposon mutant. EPA polysaccharides were extracted from three independent cultures to give average values in Fig 3C.

**S4 Figure. ^1^H-^13^C HSQC spectra showing signals altered in *epa* mutants. A.** Region corresponding to anomeric protons (4.2-5.5 ppm) and anomeric carbons (90-105 ppm) highlighting four regions of the spectra (boxed) with signals shifted or changing in intensity in the *epa* mutants. **B.** Boxed regions in A. are shown for individual mutant and one complemented strain.

**S5 Figure. PCR analysis of OG1RF derivatives harbouring in-frame deletions in *OG1RF_11720, OG1RF_11715, OG1RF_11714* and *OG1RF_11707*.**

**S6 Figure. Virulence of *epa* transposon mutants and complemented strains in the zebrafish model of infection.** Survival of zebrafish larvae (n>20) following infection with *E. faecalis* OG1RF (WT) and *epa* insertion mutant was monitored over 90h post infection. **A.** Mutant *OPDV_11720::Tn2.5*. **B.** Mutant *OPDV_11715::Tn2.13*. **C.** Mutant *OPDV_11714::Tn2.14*. **D.** Mutant *OPDV_11707::Tn2.8*. Statistical significance was determined by Log-rank test; NS, *P*>0.05; \*\**P*<0.01; *** *P*<0.001; **** *P*<0.0001. The same data corresponding to the WT strain are shown in Fig4A/4C and Fig 4B/4D.

**S7 Figure. Comparative analysis of OG1RF and *OPDV* virulence in the zebrafish model of infection.** Survival of zebrafish larvae (n=28) following infection with 1,000 CFUs of *E. faecalis* OG1RF (WT) and *OPDV* mutant was monitored over 90 h post infection. The lack of statistical significance (P=0.645) was determined by Log-rank test.

**S8 Figure. Growth rate analysis of *E. faecalis* OG1RF and *epa* derivatives.** Cells from overnight cultures in BHI were diluted to an OD at 600 nm of 0.01 in 25 ml BHI and growth of standing cultures was monitored over 6 h. The data presented are the average of 3 independent cultures. The same OG1RF growth curves were used as a control in each graph.

**S9 Figure. Step-by-step description of the transposon mapping strategy. A.** Schematic representation of the *mariner* transposon used. It consists of a gentamycin resistance cassette flanked by two inverted repeats. **B.** Step 1: digestion of chromosomal DNA with SspI, which has a unique cleavage site in the gentamycin resistance cassette. **C.** step 2: self-ligation of SspI digestion products. **D.** step 3: reverse PCR on ligation products with two divergent oligonucleotides (Mar_dn and Mar_up). **E.** step 4: sequencing of the PCR product using oligonucleotide T7.

**S1 Table. Electrophoretic mobility measurements.** The values presented are the average of three independent biological replicates ± standard deviation.

**S2 Table. Statistical significance of pairwise comparisons of electrophoretic mobility.** The significance values have been calculated using two-way ANOVA.

**S3 Table. Bacterial strains, plasmids and oligonucleotides used in this study.**

