## Supplementary material for "Decoration of the enterococcal polysaccharide antigen EPA is essential for virulence, cell surface charge and resistance to innate immunity"

### Slide 1
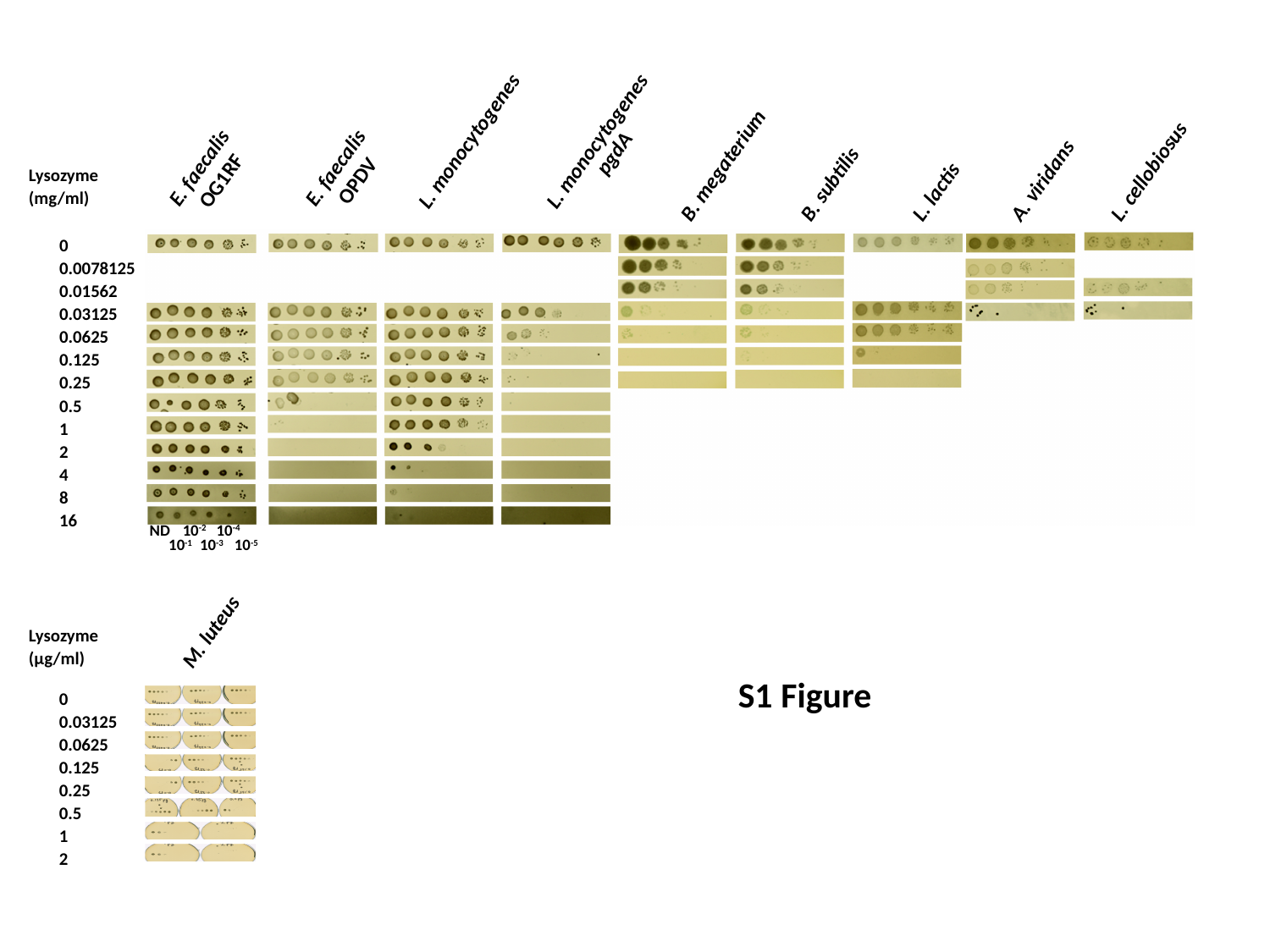

L. monocytogenes
L. monocytogenes
pgdA
E. faecalis
OG1RF
E. faecalis
OPDV
B. megaterium
L. cellobiosus
Lysozyme
(mg/ml)
A. viridans
B. subtilis
L. lactis
0
0.0078125
0.01562
0.03125
0.0625
0.125
0.25
0.5
1
2
4
8
16
ND 10^-2^ 110^-4^
10^-1^ 10^-3-1^ 10^-5^
M. luteus
Lysozyme
(µg/ml)
S1 Figure
0
0.03125
0.0625
0.125
0.25
0.5
1
2
