## Supplementary figures and images for "Decoration of the enterococcal polysaccharide antigen EPA is essential for virulence, cell surface charge and resistance to innate immunity"

### Supplementary file 2

#### Slide 1
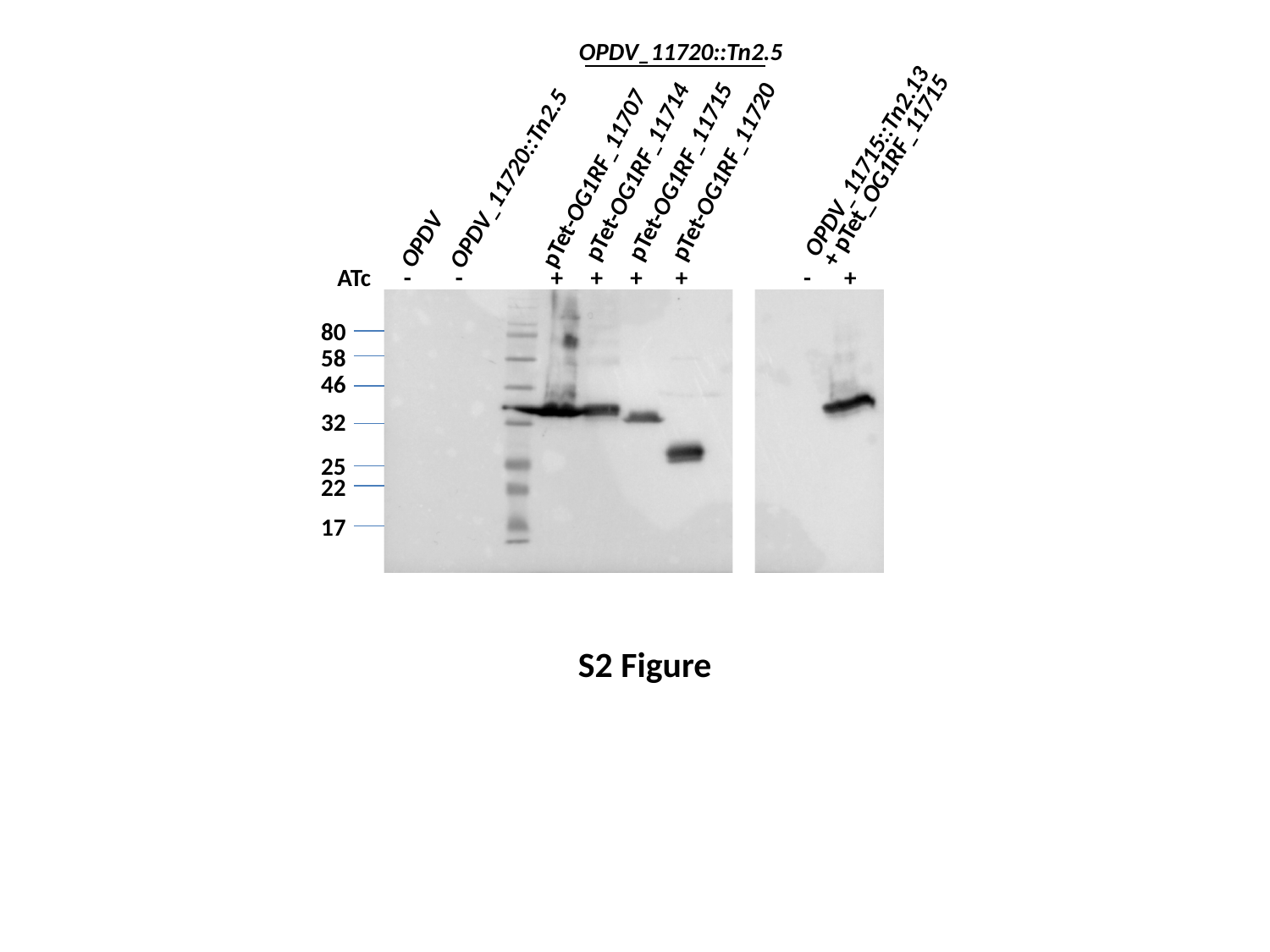

OPDV_11720::Tn2.5
OPDV_11715::Tn2.13
+ pTet_OG1RF_11715
pTet-OG1RF_11714
pTet-OG1RF_11715
pTet-OG1RF_11720
pTet-OG1RF_11707
OPDV_11720::Tn2.5
OPDV
ATc - - + + + +
 - +
80
58
46
32
25
22
17
S2 Figure

### Supplementary file 4

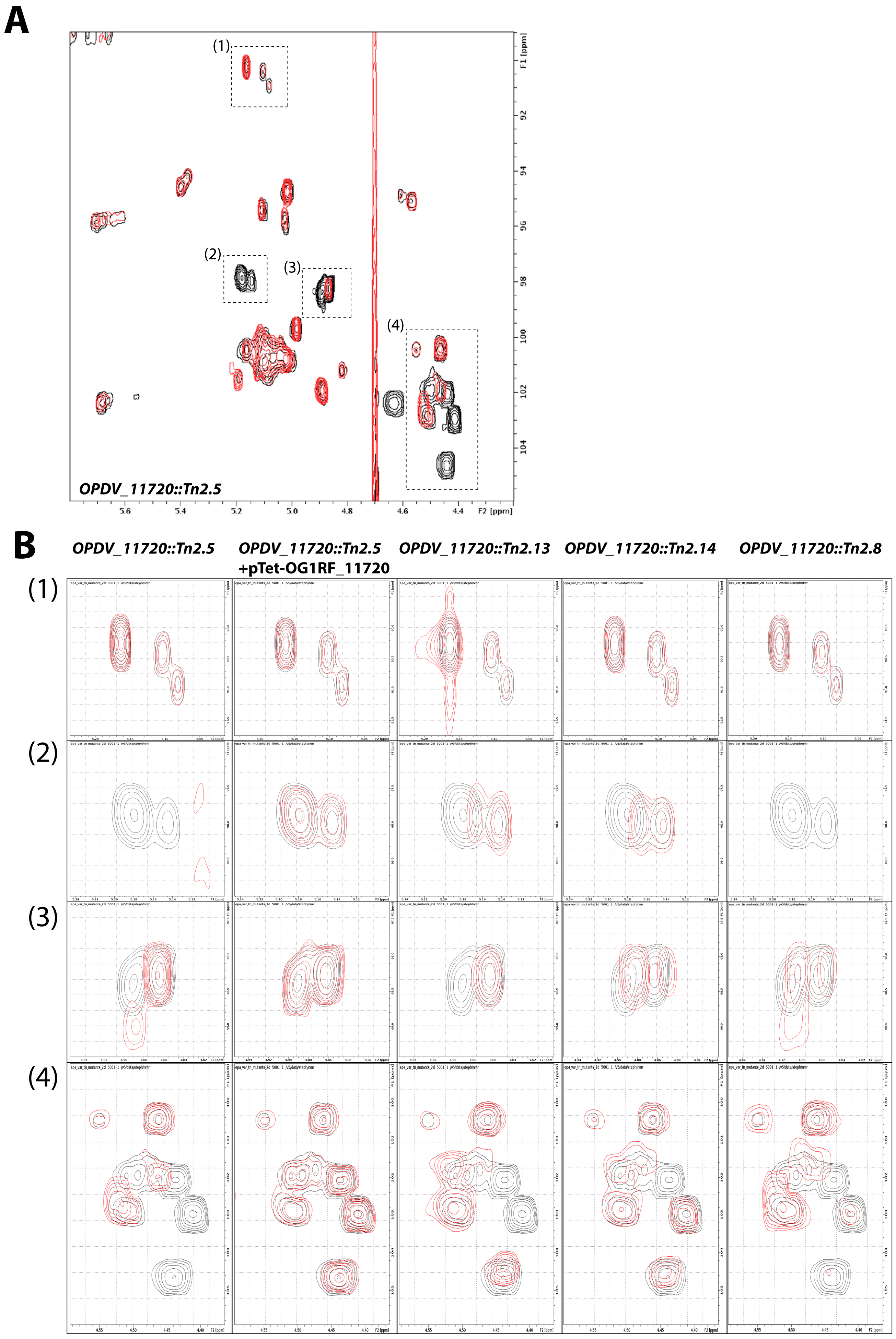

### Supplementary file 7

#### Slide 1
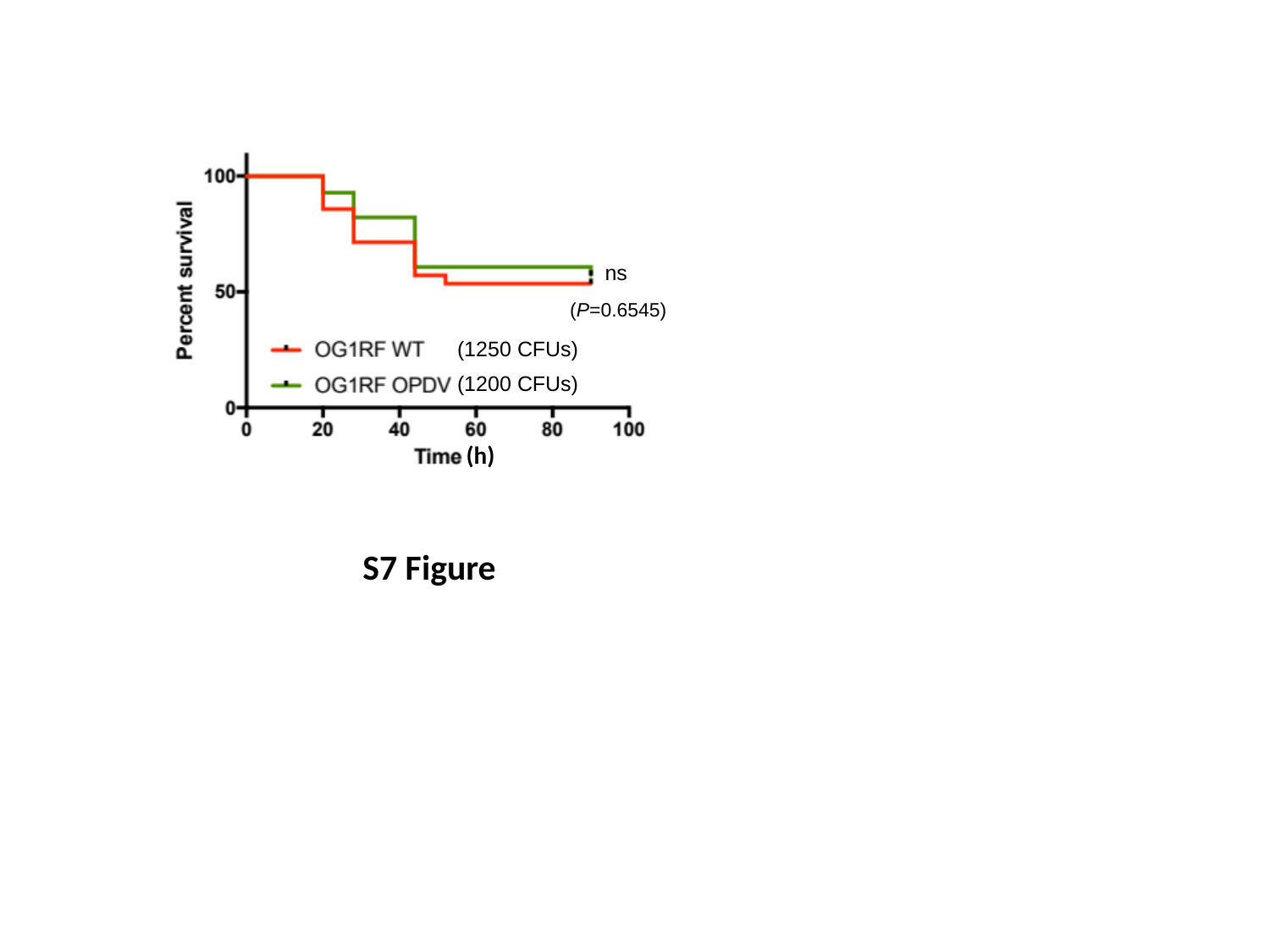

ns
(P=0.6545)
(1250 CFUs)
(1200 CFUs)
(h)
S7 Figure
