## Supplementary material for "Decoration of the enterococcal polysaccharide antigen EPA is essential for virulence, cell surface charge and resistance to innate immunity"

### Slide 1
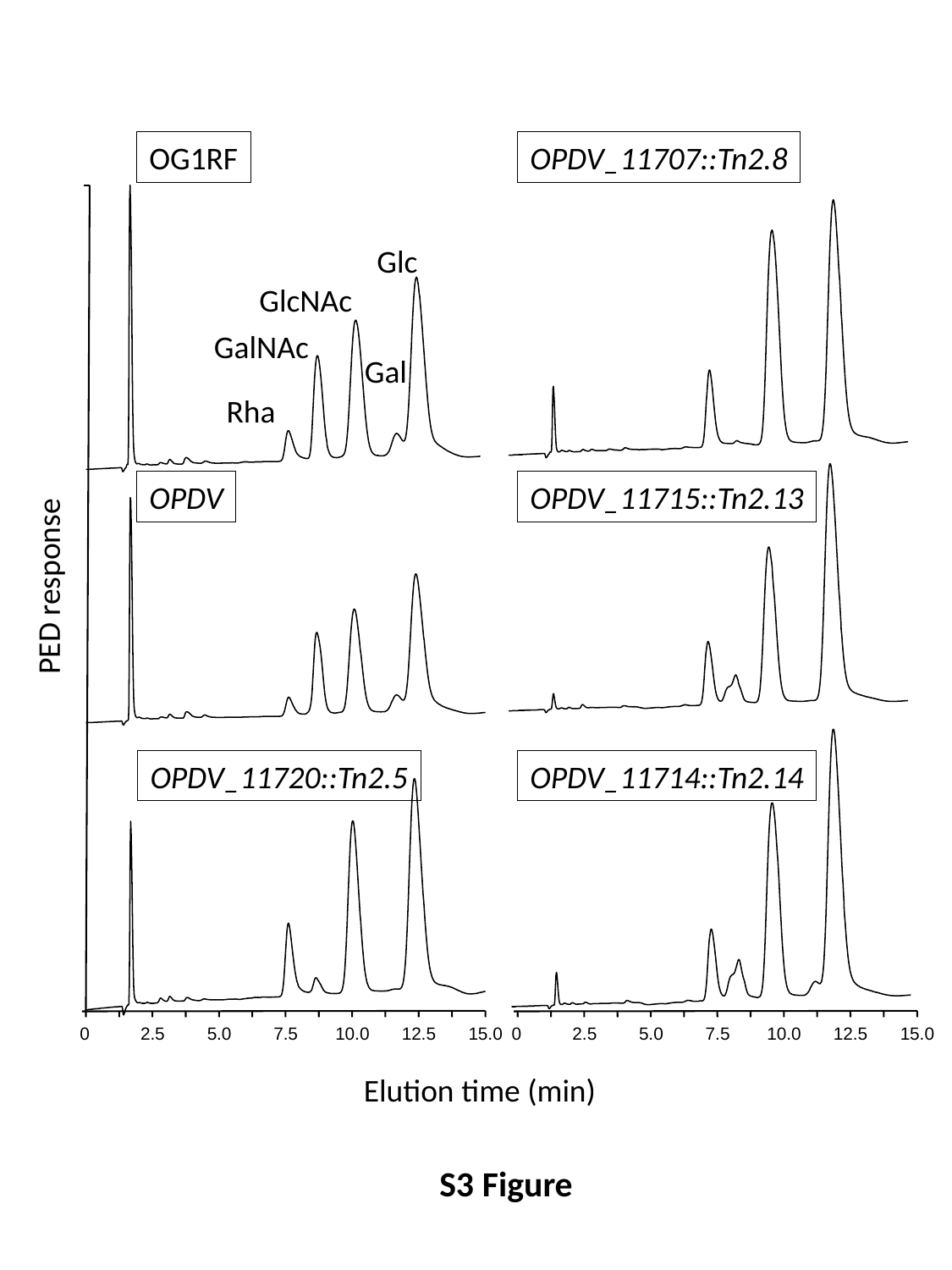

OG1RF
OPDV_11707::Tn2.8
Glc
GlcNAc
GalNAc
Gal
Rha
OPDV
OPDV_11715::Tn2.13
PED response
OPDV_11720::Tn2.5
OPDV_11714::Tn2.14
0
2.5
5.0
7.5
10.0
12.5
15.0
0
2.5
5.0
7.5
10.0
12.5
15.0
Elution time (min)
S3 Figure
