## Supplementary material for "Decoration of the enterococcal polysaccharide antigen EPA is essential for virulence, cell surface charge and resistance to innate immunity"

### Slide 1
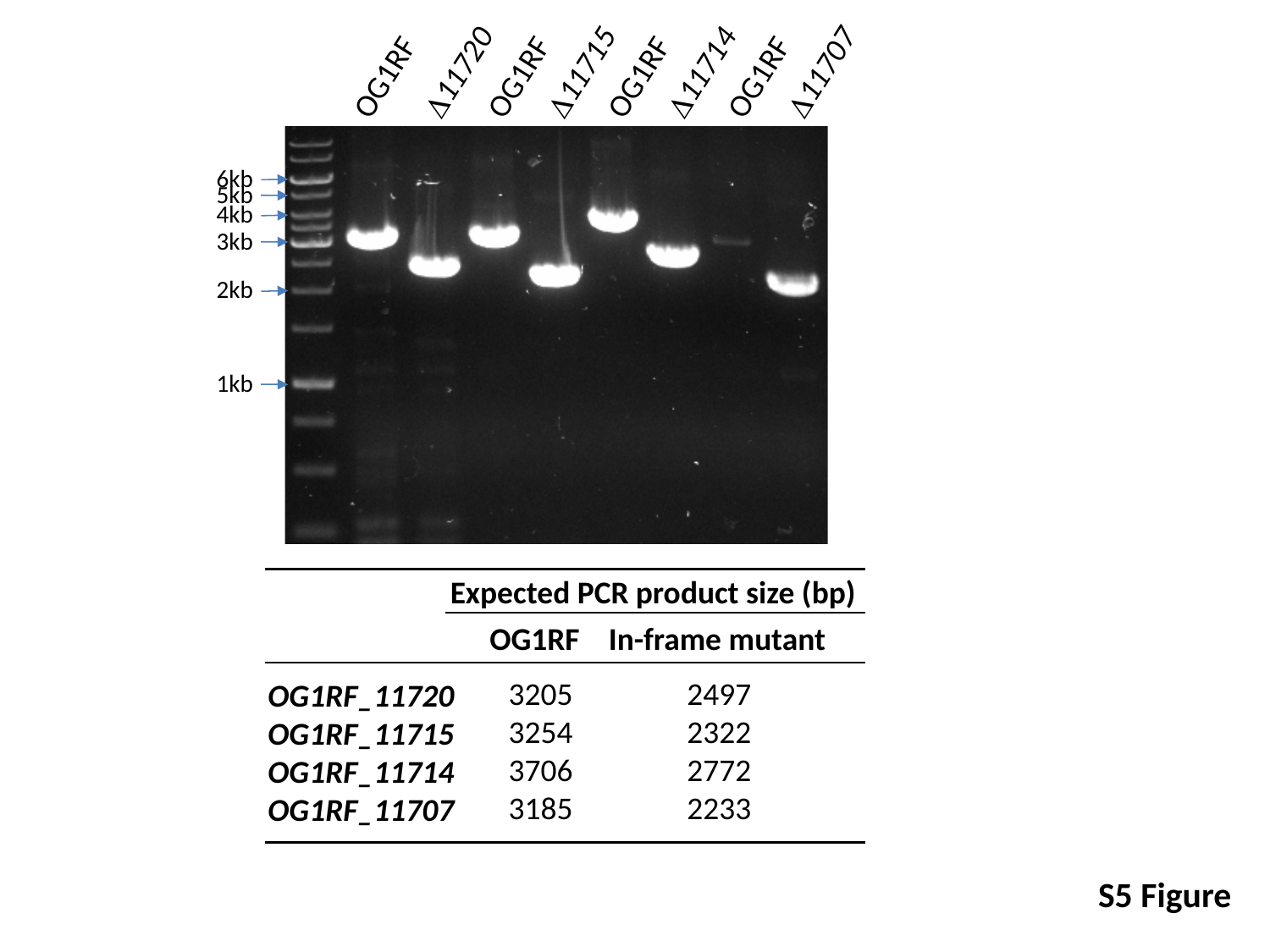

Δ11720
Δ11715
Δ11714
Δ11707
OG1RF
OG1RF
OG1RF
OG1RF
6kb
5kb
4kb
3kb
2kb
1kb
Expected PCR product size (bp)
OG1RF
In-frame mutant
3205
3254
3706
3185
2497
2322
2772
2233
OG1RF_11720
OG1RF_11715
OG1RF_11714
OG1RF_11707
S5 Figure
