## Supplementary material for "Decoration of the enterococcal polysaccharide antigen EPA is essential for virulence, cell surface charge and resistance to innate immunity"

### Slide 1
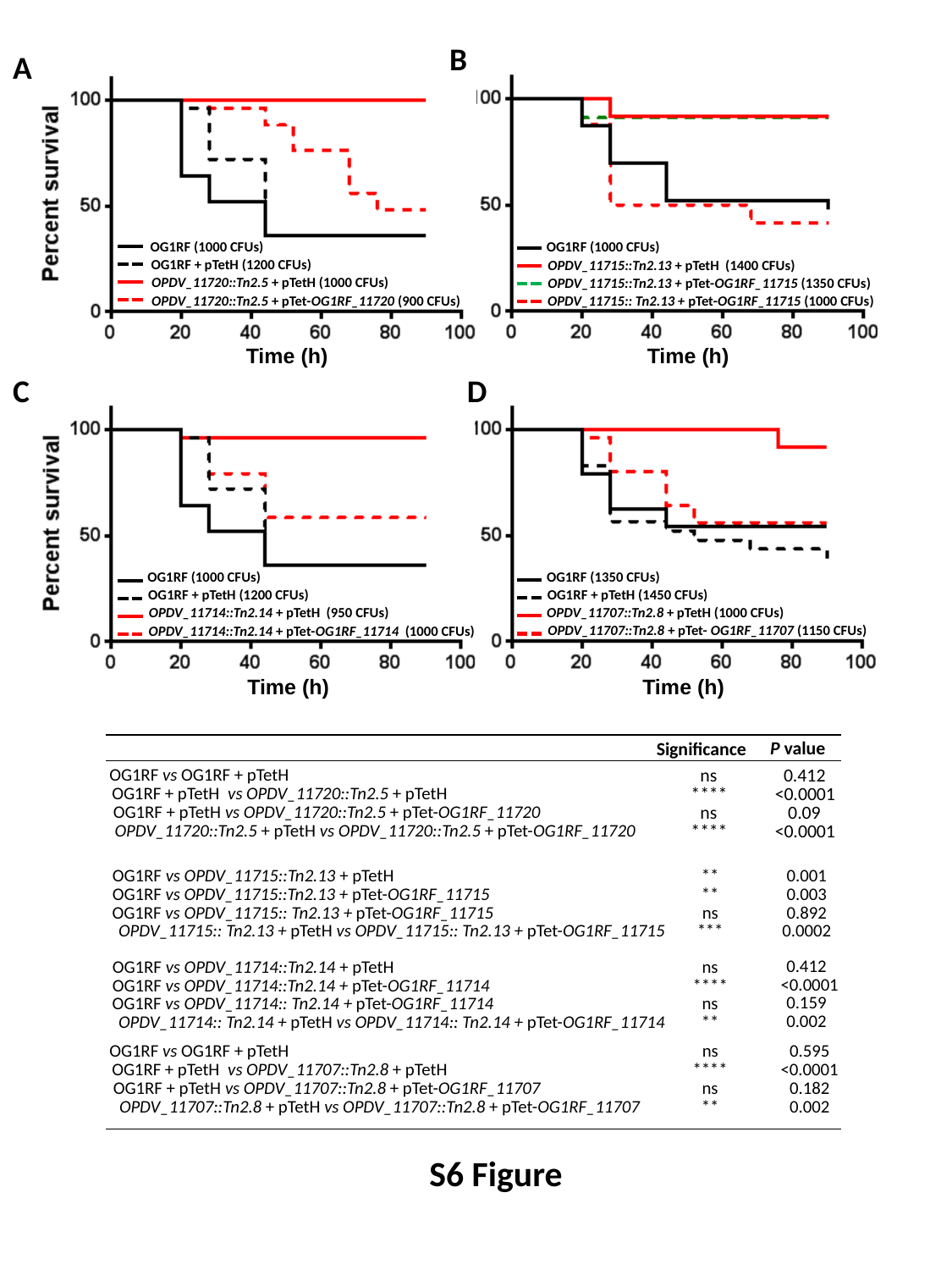

B
A
OG1RF (1000 CFUs)
OG1RF (1000 CFUs)
OPDV_11715::Tn2.13 + pTetH (1400 CFUs)
OG1RF + pTetH (1200 CFUs)
OPDV_11715::Tn2.13 + pTet-OG1RF_11715 (1350 CFUs)
OPDV_11720::Tn2.5 + pTetH (1000 CFUs)
OPDV_11715:: Tn2.13 + pTet-OG1RF_11715 (1000 CFUs)
OPDV_11720::Tn2.5 + pTet-OG1RF_11720 (900 CFUs)
Time (h)
Time (h)
C
D
OG1RF (1350 CFUs)
OG1RF (1000 CFUs)
OG1RF + pTetH (1450 CFUs)
OG1RF + pTetH (1200 CFUs)
OPDV_11707::Tn2.8 + pTetH (1000 CFUs)
OPDV_11714::Tn2.14 + pTetH (950 CFUs)
OPDV_11707::Tn2.8 + pTet- OG1RF_11707 (1150 CFUs)
OPDV_11714::Tn2.14 + pTet-OG1RF_11714 (1000 CFUs)
Time (h)
Time (h)
P value
0.412
<0.0001
0.09
<0.0001
Significance
ns
****
ns
****
OG1RF vs OG1RF + pTetH
OG1RF + pTetH vs OPDV_11720::Tn2.5 + pTetH
OG1RF + pTetH vs OPDV_11720::Tn2.5 + pTet-OG1RF_11720
OPDV_11720::Tn2.5 + pTetH vs OPDV_11720::Tn2.5 + pTet-OG1RF_11720
**
**
ns
***
0.001
0.003
0.892
0.0002
OG1RF vs OPDV_11715::Tn2.13 + pTetH
OG1RF vs OPDV_11715::Tn2.13 + pTet-OG1RF_11715
OG1RF vs OPDV_11715:: Tn2.13 + pTet-OG1RF_11715
OPDV_11715:: Tn2.13 + pTetH vs OPDV_11715:: Tn2.13 + pTet-OG1RF_11715
0.412
<0.0001
0.159
0.002
ns
****
ns
**
OG1RF vs OPDV_11714::Tn2.14 + pTetH
OG1RF vs OPDV_11714::Tn2.14 + pTet-OG1RF_11714
OG1RF vs OPDV_11714:: Tn2.14 + pTet-OG1RF_11714
OPDV_11714:: Tn2.14 + pTetH vs OPDV_11714:: Tn2.14 + pTet-OG1RF_11714
OG1RF vs OG1RF + pTetH
ns
****
ns
**
0.595
<0.0001
0.182
0.002
OG1RF + pTetH vs OPDV_11707::Tn2.8 + pTetH
OG1RF + pTetH vs OPDV_11707::Tn2.8 + pTet-OG1RF_11707
OPDV_11707::Tn2.8 + pTetH vs OPDV_11707::Tn2.8 + pTet-OG1RF_11707
S6 Figure
