## Supplementary material for "Decoration of the enterococcal polysaccharide antigen EPA is essential for virulence, cell surface charge and resistance to innate immunity"

### Slide 1
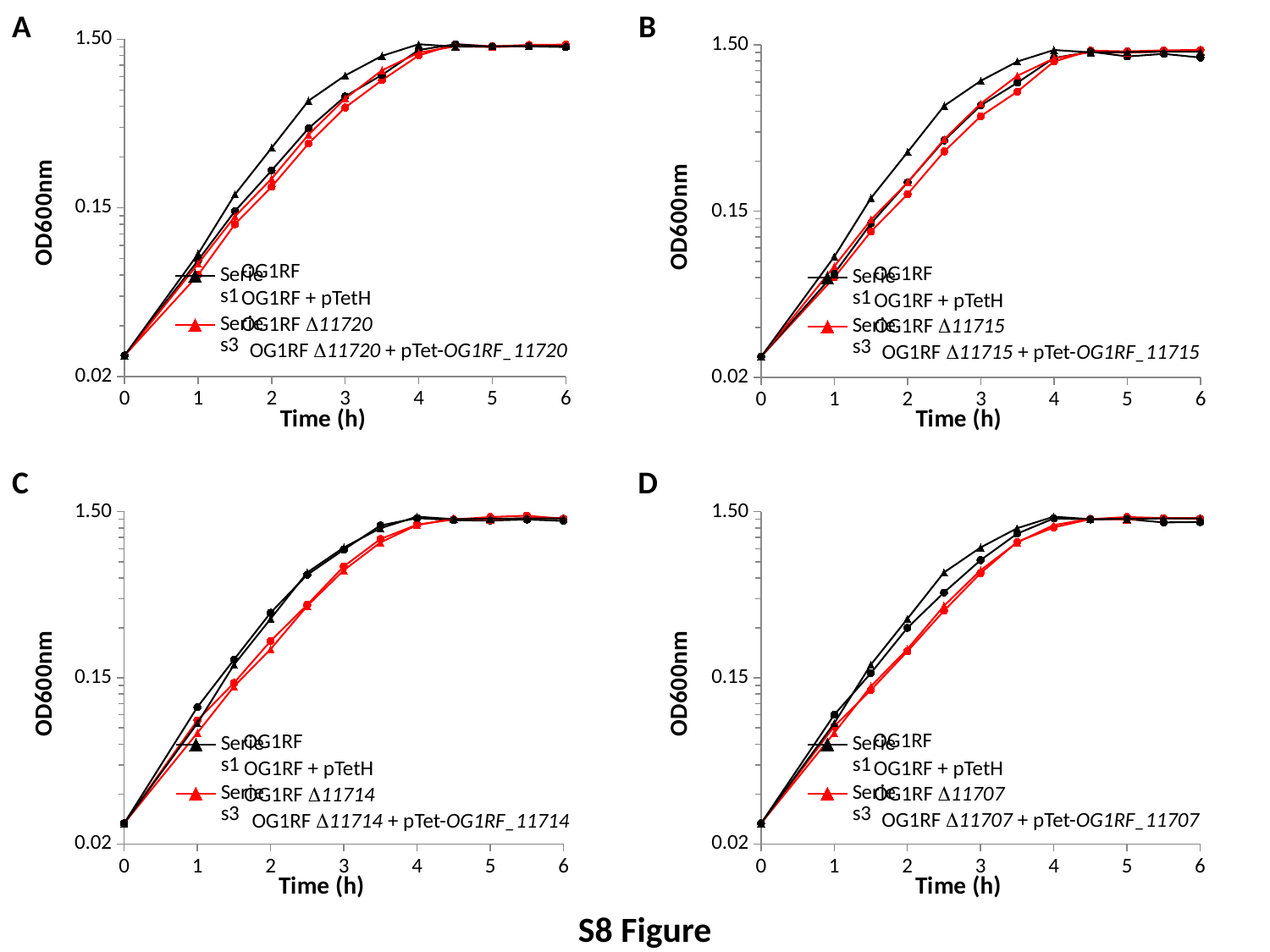

A
B
#### Chart
| Category |
|---|
#### Chart
| Category | | | | |
|---|---|---|---|---|OG1RF
OG1RF
OG1RF + pTetH
OG1RF + pTetH
OG1RF Δ11720
OG1RF Δ11715
OG1RF Δ11720 + pTet-OG1RF_11720
OG1RF Δ11715 + pTet-OG1RF_11715
C
D
#### Chart
| Category |
|---|
#### Chart
| Category | | | | |
|---|---|---|---|---|OG1RF
OG1RF
OG1RF + pTetH
OG1RF + pTetH
OG1RF Δ11707
OG1RF Δ11714
OG1RF Δ11707 + pTet-OG1RF_11707
OG1RF Δ11714 + pTet-OG1RF_11714
S8 Figure
