## Supplementary material for "Decoration of the enterococcal polysaccharide antigen EPA is essential for virulence, cell surface charge and resistance to innate immunity"

### Slide 1
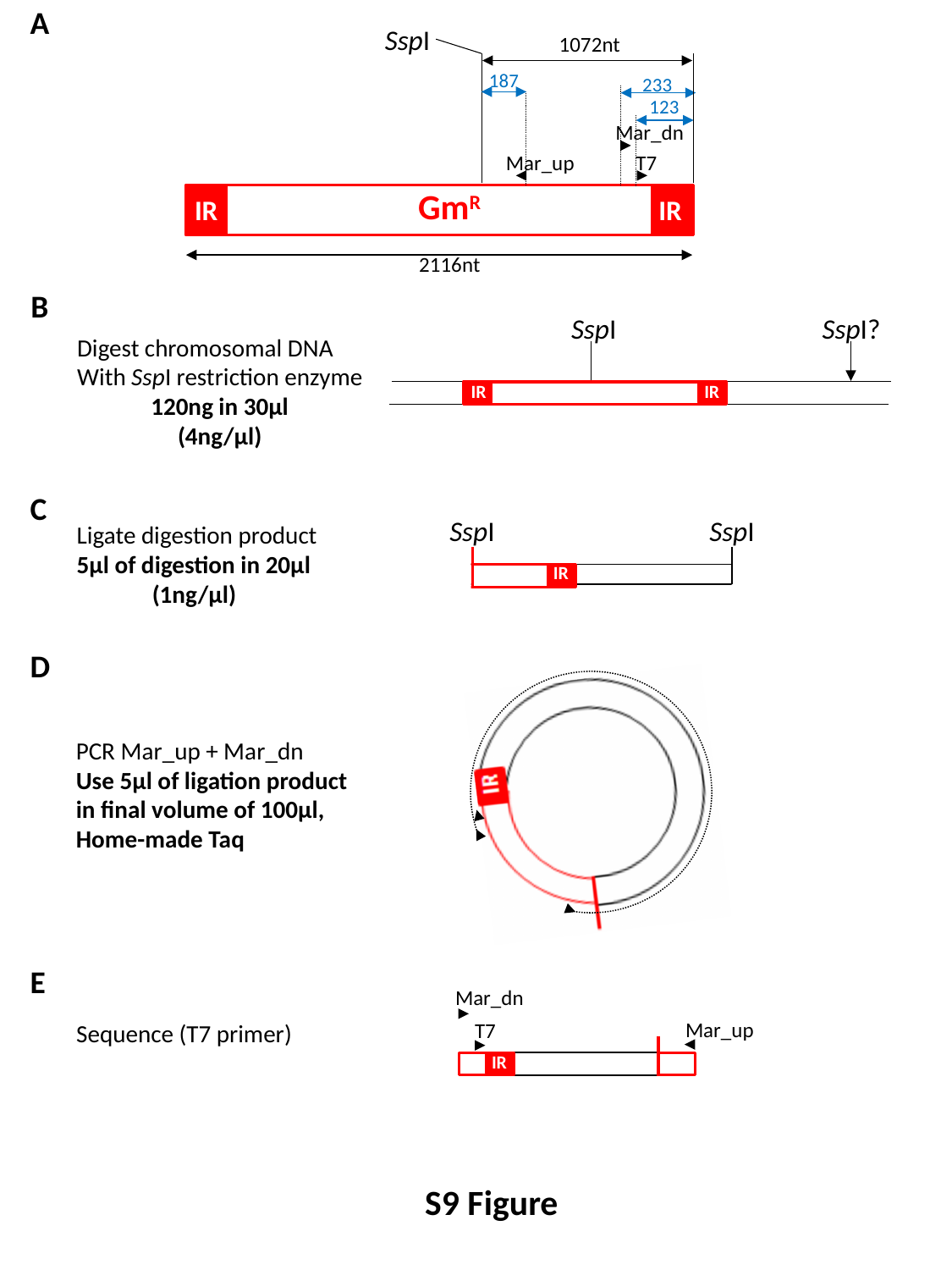

A
SspI
1072nt
233
187
123
Mar_dn
Mar_up
T7
Gm^R^
IR
IR
2116nt
B
SspI
SspI?
Digest chromosomal DNA
With SspI restriction enzyme
120ng in 30µl
(4ng/µl)
IR
IR
C
SspI
SspI
Ligate digestion product
5µl of digestion in 20µl
(1ng/µl)
D
PCR Mar_up + Mar_dn
Use 5µl of ligation product
in final volume of 100µl,
Home-made Taq
E
Mar_dn
Sequence (T7 primer)
Mar_up
T7
IR
S9 Figure
