## Supplementary material for "Decoration of the enterococcal polysaccharide antigen EPA is essential for virulence, cell surface charge and resistance to innate immunity"

**S1 Table. Electrophoretic mobility measurements ($10^{-8}$ $m^2V^{-1}s^{-1}$).** The values presented are the average of three independent biological replicates ± standard deviation. Each electrophoretic mobility value was measured from an average of 200 cells.

| | pH | | | | |
|---|---|---|---|---|---|
| **Strains** | **2.0** | **3.0** | **4.0** | **4.5** | **5.5** |
| **OG1RF** | -2.11 ± 0.76 | -3.29 ± 0.29 | -3.48 ± 0.24 | -3.22 ± 0.17 | -2.65 ± 0.11 |
| ***OPDV*** | -2.30 ± 0.12 | -3.27 ± 0.06 | -3.34 ± 0.13 | -3.20 ± 0.13 | -2.78 ± 0.10 |
| ***OPDV_11720::Tn2.5*** | 0.29 ± 0.07 | 0.01 ± 0.13 | -0.80 ± 0.12 | -1.00 ± 0.19 | -1.49 ± 0.13 |
| ***OPDV_11720::Tn2.5* + pTetH-*OG1RF_11720*** | -1.51 ± 0.52 | -2.91 ± 0.27 | -3.44 ± 0.26 | -3.29 ± 0.18 | -2.67 ± 0.06 |
| ***OPDV_11707::Tn2.8*** | 0.18 ± 0.08 | -0.38 ± 0.07 | -1.27 ± 0.28 | -1.51 ± 0.09 | -1.88 ± 0.23 |
| ***OPDV_11707::Tn2.8* + pTetH-*OG1RF_11707*** | -2.22 ± 0.17 | -3.26 ± 0.11 | -3.30 ± 0.18 | -3.17 ± 0.13 | -2.74 ± 0.08 |
| ***OPDV_11715::Tn2.13* (*epaOX*)** | 0.17 ± 0.05 | -0.19 ± 0.08 | -0.90 ± 0.01 | -1.14 ± 0.10 | -1.49 ± 0.07 |
| ***OPDV_11715::Tn2.13* + pTetH-*OG1RF_11715*** | -2.28 ± 0.06 | -3.15 ± 0.08 | -3.27 ± 0.17 | -3.17 ± 0.20 | -2.68 ± 0.08 |
| ***OPDV_11714::Tn2.14* (*epaX*)** | -1.05 ± 0.13 | -3.02 ± 0.02 | -3.55 ± 0.18 | -3.26 ± 0.30 | -2.81 ± 0.18 |
| ***OPDV_11714::Tn2.14* + pTetH-*OG1RF_11714*** | -2.27 ± 0.13 | -3.07 ± 0.16 | -3.10 ± 0.18 | -3.07 ± 0.11 | -2.65 ± 0.13 |

**S2 Table. Statistical significance of pairwise comparisons of electrophoretic mobility with OPDV.** The significance values have been calculated using two-way ANOVA.

| | pH | | | | |
|---|---|---|---|---|---|
| **Strains** | **2.0** | **3.0** | **4.0** | **4.5** | **5.5** |
| **OG1RF** | NS | NS | NS | NS | NS |
| ***OPDV_11720::Tn2.5*** | *** | *** | *** | *** | *** |
| ***OPDV_11720::Tn2.5* + pTetH-*OG1RF_11720*** | NS | NS | NS | NS | NS |
| ***OPDV_11707::Tn2.8*** | *** | *** | *** | *** | ** |
| ***OPDV_11707::Tn2.8* + pTetH-*OG1RF_11707*** | NS | NS | NS | NS | NS |
| ***OPDV_11715::Tn2.13* (*epaOX*)** | *** | *** | *** | *** | *** |
| ***OPDV_11715::Tn2.13* + pTetH-*OG1RF_11715*** | NS | NS | NS | NS | NS |
| ***OPDV_11714::Tn2.14* (*epaX*)** | *** | ** | NS | NS | NS |
| ***OPDV_11714::Tn2.14* + pTetH-*OG1RF_11714*** | NS | NS | NS | NS | NS |

NS, Not significant; ***, $P<0.001$; **, $P<0.01$

### S4 Table. Bacterial strains, plasmids and oligonucleotides used in this study.

| Strains, plasmids | Relevant properties or genotype[a] | Source or reference |
|---|---|---|
| **Strains** | | |
| ***Aerococcus viridans*** | | |
| ATCC11563 | | [1] |
| ***Bacillus megaterium*** | | |
| KM | | [2] |
| ***Bacillus subtilis*** | | |
| 168 | | [3] |
| ***Enterococcus faecalis*** | | |
| OG1RF | Plasmid-free, virulent strain isolated from the oral cavity | [4] |
| OG1RF OPDV | OG1RF derivative with deletions in *oatA*, *pgdA*, *dltA* and *sigV* | This work |
| ***Enterococcus faecium*** | | |
| DO (TX16) | | [5] |
| ***Enterococcus hirae*** | | |
| ATCC9790 | | [6] |
| ***Escherichia coli*** | | |
| TG1 | Host for plasmid propagation | NEB |
| TG1(*RepA*) | TG1 derivative harboring *RepA* for pGhost propagation at 37°C | P. Serror |
| ***Listeria monocytogenes*** | | |
| EGDe | | [7] |
| EGDe $\Delta$*pgdA* | | [7] |
| ***Streptococcus agalactiae*** | | |
| NEM316 | | [8] |
| ***Streptococcus gordonii*** | | |
| DL-1 Challis | Previously called *Streptococcus sanguis* | [9] |
| ***Streptococcus mutans*** | | |
| UA159 | | [10] |
| ***Streptococcus gallolyticus*** | | |
| UCN34 | | [11] |
| **Plasmids** | | |
| pTetH | pAT18 derivative for tetracycline-inducible expression in *E. faecalis* | Lab stock |
| pGhost9 | Thermosensitive plasmid used for gene replacement | [12] |
| pGHH_11707 | pGhost9 derivative used for in-frame deletion of *OG1RF_11707* | This work |
| pGHH_11714 | pGhost9 derivative used for in-frame deletion of *OG1RF_11714* | This work |
| pGHH_11715 | pGhost9 derivative used for in-frame deletion of *OG1RF_11715* | This work |
| pGHH_11720 | pGhost9 derivative used for in-frame deletion of *OG1RF_11720* | This work |
| pMAD-$\Delta$*oatA* | pMAD derivative used to build the in-frame *oatA* deletion | [13] |
| pMAD-$\Delta$*pgdA* | pMAD derivative used to build the in-frame *pgdA* deletion | [13] |
| pMAD-$\Delta$*dltA* | pMAD derivative used to build the in-frame *dltA* deletion | [14] |
| pMAD-$\Delta$*sigV* | pMAD derivative used to build the in-frame *sigV* deletion | [14] |
| pTet-*OGR1F_11720* | pTetH derivative encoding *OG1RF_11720* | This work |
| pTet-*OGR1F_11715* | pTetH derivative encoding *OG1RF_11715* | This work |
| pTet-*OGR1F_11714* | pTetH derivative encoding *OG1RF_11714* | This work |
| pTet-*OGR1F_11707* | pTetH derivative encoding *OG1RF_11707* | This work |
| **Oligonucleotides[a]** | | |
| 11720_Fw | GGGCCATGGAAACAGCACTTGTTTCAATTATTATGC | |
| 11720_Rev | TTTGGATCCTTCATTCTTTGCATATTTAAATGTTGTAT | |
| 11715_Fw | CCCCCATGGAAAAAGAAAATTTAAAGTTAAGCGTGATTATTCC | |
| 11715_Rev | TATGGATCCCTTCCCTAAAATTTTTCGATACATAAAATTATATA | |
| 11714_Fw | AAACCATGGAAATTAGTATAATTGTTCCTGTTT | |
| 11714_Rev | CCCGGATCCCTTATTAACCTTTCTTAAATACCTCTGTTGCA | |
| 11707_Fw | AAACCATGGAAACATTCCTAATCACAGGCGGC | |
| 11707_Rev | GGGGGATCCTTCTTTTAATTCATAATTAACATATTTATCCAAAC | |

| | |
|---|---|
| 11720_H11 | CTTTGGCGAATTCTTGTTTATTATCTTTACTTAAAGGTC |
| 11720_H12 | CTATTCATTCTTTGCATACATAATAATTGAAACAAGTGCTGTTTCC |
| 11720_H21 | CAATTATTATGTATGCAAAGAATGAATAGGGAGGAAAAGGAAAAG |
| 11720_H22 | AAACTCGAGGACATACATACTATAGTGATATGCTGAAACTTTTAAG |
| 11715_H11 | AAACTCGAGAAGTCAAAGAGGCTATGTTCGGTCATC |
| 11715_H12 | CTTCCCTAACACATTATATACTGGAATAATCACGCTTAAC |
| 11715_H21 | ATTCCAGTATATAATGTGTTAGGGAAGTAAGGAGTTAAAGATGTCAG |
| 11715_H22 | AAAGGTACCCTTATTAAATCCACCTCTCAGTATAGCTAC |
| 11714_H11 | AAAGAATTCCATTTTGTACATGTTAAGTTCCTTGAACATG |
| 11714_H12 | CCTCTGTTGCATAACAGGAACAATTATACTAATTTCTGACATC |
| 11714_H21 | AATTGTTCCTGTTATGCAACAGAGGTATTTAAGAAAGGTTAATAAG |
| 11714_H22 | AAACTCGAGCTAAGCAACTCTTTTTCTGTTAAAGCAAAC |
| 11707_H11 | AAAGAATTCGAGCTGATCATTCTATAGAAAGATGTACTTATG |
| 11707_H12 | GGACTGAATACTGCCGCCTGTGATTAGGAATGTTTCCAC |
| 11707_H21 | CTAATCACAGGCGGCAGTATTCAGTCCGGTTTGGATAAATATG |
| 11707_H22 | AAAGGTACCGTTTCTTAGTCTCTAAAAATACACGGCCAAC |

[a] restriction sites used for cloning are underlined (CCATGG, NcoI; GGATCC, BamHI; GAATTC, EcoRI; GGTACC, KpnI; CTCGAG, XhoI)
